# Precise genotyping of circular mobile elements uncovers human associated plasmids with surprisingly recent common ancestors

**DOI:** 10.1101/2021.05.25.445656

**Authors:** Nitan Shalon, David Relman, Eitan Yaffe

## Abstract

Mobile genetic elements with circular genomes play a key role in the evolution of microbial communities. These circular genomes correspond to cyclic paths in metagenome graphs, and yet, assemblies derived from natural microbial communities produce graphs riddled with spurious cycles, complicating the accurate reconstruction of circular genomes. We present an algorithm that reconstructs true circular genomes based on the identification of so-called ‘dominant’ cycles. Our algorithm leverages paired reads to bridge gaps between assembly contigs and scrutinizes cycles through a nucleotide-level analysis, making the approach robust to mis-assembly artifacts. We validated the approach using simulated and reference data. Application of this approach to 32 publicly available DNA shotgun sequence data sets from diverse natural environments led to the reconstruction of hundreds of circular mobile genomes. Clustering revealed 20 clusters of cryptic, prevalent, and abundant plasmids that have clonal population structures with surprisingly recent common ancestors. This work enables the robust study of evolution and spread of mobile elements in natural settings.

## INTRODUCTION

Horizontal gene transfer (HGT) is a major driver of microbial evolution^1^. HGT supports the rapid adaptation of microbes to ecological niches^2,3^ and can facilitate the spread of virulence factors and antimicrobial resistance determinants within and between microbial species^4–6^. Extrachromosomal circular mobile genetic elements (ecMGEs), such as plasmids and phage with circular genomes, are of particular interest since they are potential intermediates of HGT^7^. Experimental methods have been developed to enrich for ecMGEs from complex microbial communities, including the physical enrichment for plasmid^8^ or viral particles^9^, or the removal of linear DNA followed by multiple displacement amplification^10^. Due to reduced cost and benchwork simplicity, MGE characterization is shifting towards metagenomic, or shotgun sequencing, requiring the development of robust computational tools that reliably infer MGEs from metagenomic data^11,12^. While numerous computational tools have been developed to identify MGE-associated sequences from metagenomic data^11–16^, they typically rely on reference sequences and can conflate extrachromosomal with integrated forms of MGE. Moreover, reference-based approaches classify only sequence fragments (known as ‘contigs’) and do not reliably recover near-complete genomes. Having near-complete MGE genomes is instrumental to the study of HGT dynamics, such as for tracking plasmids bearing antimicrobial resistance determinants in hospital settings^17^.

Recently, new computational approaches have been developed that work directly from metagenomic data to recover ecMGEs, circumventing reliance on incomplete and biased reference data. These approaches traverse the assembly graph, in which vertices represent contigs and edges represent contig-contig adjacencies supported by read pairs, searching for graph components that correspond to putative ecMGEs^18–20^. However, the heuristic nature of these approaches makes them susceptible to high false-positive rates when faced with complex scenarios. A recent review^21^ reported that the precision of these reference-free tools is below 0.75. The recovery of circular genomes from the assembly graph is challenging due to repeated genetic sequences within and between genomes that produce complex graphs riddled with spurious cycles. Both plasmids^22^ and phage^23^ contribute to graph complexity since they evolve through extensive genome rearrangements, and can maintain intra-community genetic variants differing by only a few genome rearrangements^17,24,25^. A computational approach that reliably distinguishes between true circular molecules and graph artifacts is needed to genotype ecMGEs from complex communities in a robust manner. The precise recovery of near-complete MGE genomes in metagenomes will allow the field to move beyond isolate-based studies and leverage high-throughput computational surveys of mobile elements, strengthening the understanding of mobile element evolution and the emergence and dissemination of antimicrobial resistance in natural environments.

To address this gap, we have developed a new tool (“DomCycle”) for reconstructing ecMGE genomes from metagenomic data with a high degree of confidence. Our approach is based a new concept known as a “dominant cycle”, which likely correspond to true circular genomes. We present an algorithm for identifying all dominant cycles in an assembly graph, and evaluate the performance of the approach using reference data and simulated evolutionary scenarios. Application of the algorithm to data from 32 samples collected from human stool, sewage, and marine environments, revealed 20 clusters of ecMGEs that are prevalent across samples, abundant within their respective communities, and exhibit a surprisingly simple clonal population structure with little recombination and low sequence diversity.

## RESULTS

We developed a theoretical framework and associated tool (DomCycle) to recover near-complete ecMGEs from metagenome assembly graphs (**Figure 1**). In the assembly graph, *circuits* correspond to circular chains of contigs, and *cycles* are simple circuits that do not include a contig more than once. While every circular genome produces a corresponding circuit in the graph, inferring true genomes based on cycles can be challenging, as illustrated by a manufactured configuration (**Figure 1a**) and its corresponding graph (**Figure 1b**). The presence of repeat elements can result in spurious cycles (called here ‘phantom’ cycles), for which no corresponding genome exists in the biological sample. Our goal was to develop a tool that can reliably distinguish between phantom and real cycles.

**Figure 1.**
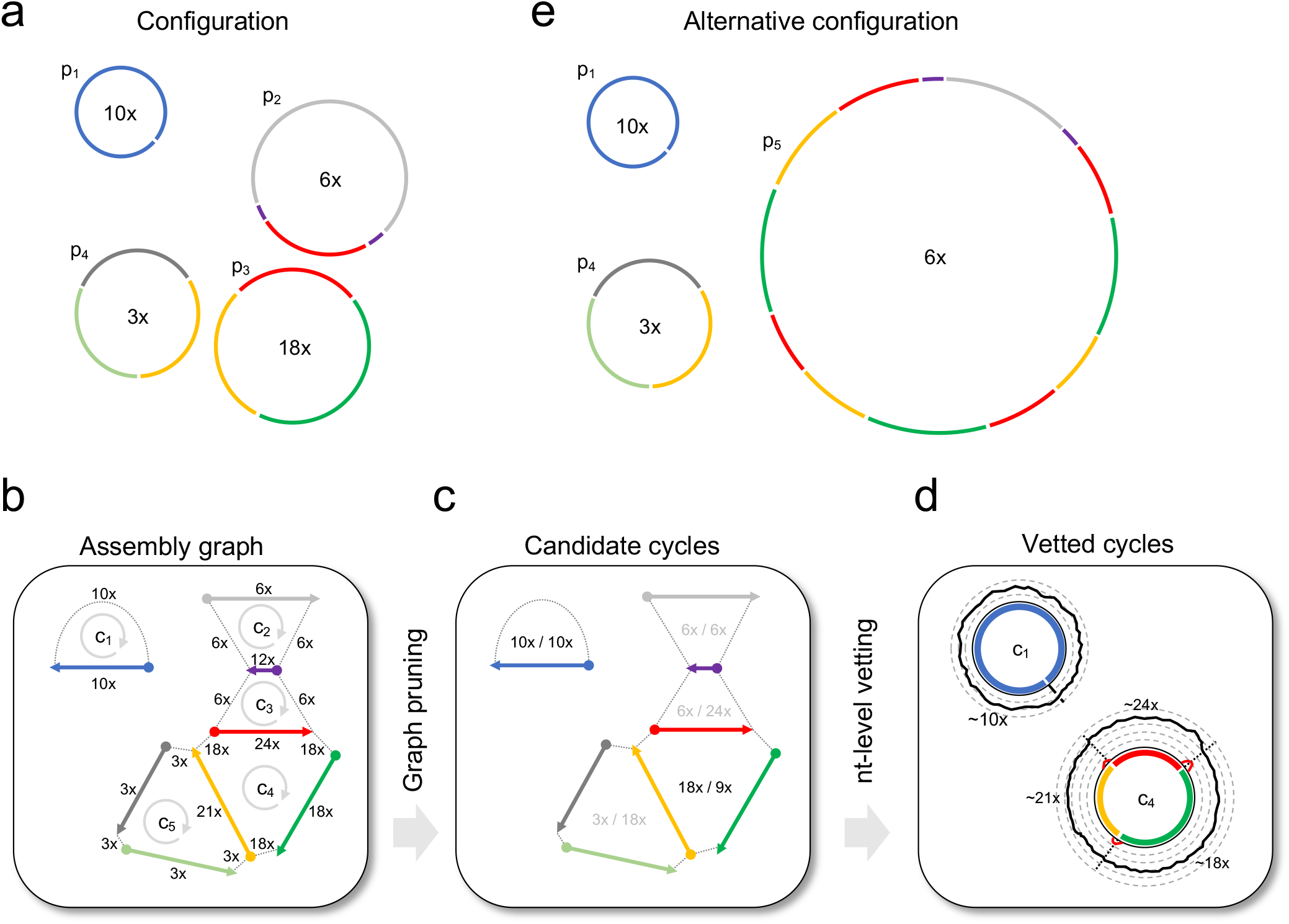
Approach overview. **a)** A manufactured example of a latent genome configuration and assembly with 8 unique contigs. Contigs are color coded and DNA genomes are marked p_1_-p_4_ with their x-coverage specified in their center. For example, p_2_ has an x-coverage of 6x and contains three unique contigs, as the short purple contig appears twice. **b)** The assembly graph of the configuration in 1a is constructed using mapped paired reads. The graph contains five cycles (c_1_-c_5_) and the x-coverage of edges is indicated. In this example, cycles c_2_ and c_3_ are phantom cycles. **c)** The cycle score σ/_c_ τ/_c_ is specified in the center of each cycle. The algorithm recovers all candidate dominant cycles (σ/_c_ τ/_c_ > 1, score colored black) and discards non-dominant cycles (score colored gray). For example, the genome p_3_ produced the cycle c_4_ which has a score of 2 (σ/_c_ = 18x, τ/_c_ = 9x), making it a candidate dominant cycle. Cycles c_2_,c_3_ are examples of phantom cycles, since each does not have a corresponding genome. **d)** Nucleotide-level read profiles are computed for all candidate dominant cycles using all paired reads for which at least one side mapped to the cycle, and the algorithm returns dominant cycles with estimates of x-coverage. Reads are grouped into cycle-supporting reads (black line) and non-supporting reads (red line). For example, the x-coverage of supporting reads along c_4_ varies, with three short stretches of non-supporting reads on contig-contig seams. Average read x-coverage values for portions of the cycles are shown on the plot. **e)** An alternative genome configuration that produces the same graph (shown in 1b). The presence of the complex genome p_5_, which corresponds to an involved circuit in the graph, affects the multiplicity of all visited cycles. For example, the multiplicity of c_4_ is equal to 3 since the circuit that corresponds to p_5_ makes on its path 3 complete turns in c_4_.

For every cycle *c*, the *bottleneck coverage σ_c_* is the minimal read coverage along the edges of the cycle, the *external coverage τ_c_* is the total number of paired reads leading in or out of the cycle (averaging in and out), and the cycle score is the ratio *σ_c_*/*τ_c_* (**Figure 1c**). We define a cycle as *dominant* if *σ_c_*/*τ_c_* > 1 and developed an algorithm that recovers all dominant cycles in the assembly graph (**Methods**). In the implementation of the algorithm, contig-contig edges in the assembly graph are inferred using paired reads while bridging possible gaps. Candidate cycles are vetted on a nucleotide-level basis, making the approach robust to common forms of mis-assembly and coverage stochasticity due to read sampling (**Figure 1d**). The algorithm yields near-complete circular genomes (near-complete and not complete, due to possible small gaps between consecutive contigs) that correspond to all vetted dominant cycles.

The latent variable of interest is the coverage *ψ_c_* of the genome associated with a dominant cycle *c*. A confounding latent variable is the multiplicity *μ_c_*, which is equal to the number of times the cycle contigs appear consecutively within the context of a larger circuit such as within a tandem repeat (see example in **Figure 1e**). Our main theoretical result is a lower bound on *ψ_c_* stating *σ_c_* − *μ_c_* · *τ_c_* < *ψ_c_*. This lower bound means that any dominant cycle is either real (i.e., *ψ_c_* > 0) or it has a multiplicity greater than the cycle score. While high-multiplicity cycles are theoretically possible, they require complex tandem structures (**Figure 1e**). To determine the degree to which dominant cycles correspond to actual circular genomes in the context of microbial communities we turn next to empirical data.

### Algorithm performance on reference data

We tested if phantom cycles (i.e., false-positives) are reported as dominant cycles using simulated chromosomal data of diverse bacterial species. DomCycle was applied to an assembly graph generated from a DNA library simulated from the chromosomal sequences of 100 strains including several conspecific strains (detailed in **Supp. Table 1**). By design, the generated assembly graph contained only phantom cycles since extra-chromosomal DNA was not included. DomCycle did not report any dominant cycles in this dataset, indicating that the likelihood of phantom cycles with bacterial genomes is rare (**Figure 2a**). We compared DomCycle to Recycler^20^, metaplasmidSPAdes^19^ and SCAPP^26^ with these data. On average, these 3 existing tools reported 93 false-positive elements per GB of assembly (**Figure 2b**), with a median element length of 2.3kb (**Figure 2c**). DomCycle stood out with zero false-positives for this dataset, highlighting the precision of the approach.

**Figure 2.**
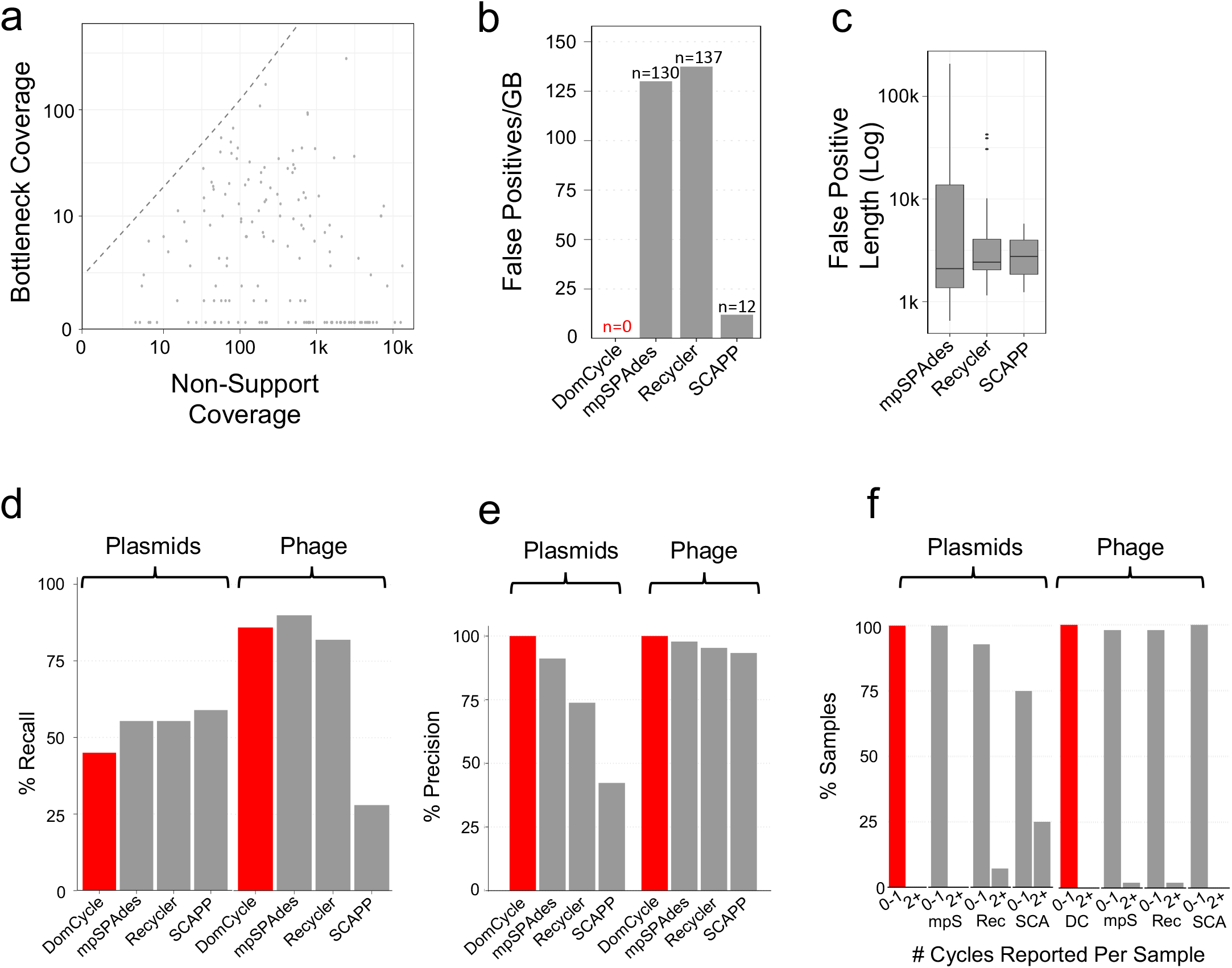
Specificity and sensitivity estimated with reference sequences. **(a)** The distribution of the bottleneck coverage vs. the total non-support coverage among cycles inspected by DomCycle on a metagenomic dataset simulated on 100 reference chromosomal sequences. The dotted line shows the threshold of significance (p = 0.01) for the global nucleotide test. All cycles are spurious (i.e., phantom) by construction and no dominant cycles were reported. **(b)** The rate of phantom cycles reported per GB of assembly on the simulated metagenomic dataset, comparing the performance of DomCycle, metaplasmidSPAdes, Recycler and SCAPP. **(c)** The length distribution for phantom cycles reported on the simulated metagenomic dataset. **(d)** Recall estimates for DomCycle, metaplasmidSPAdes, Recycler, and SCAPP on 56 reference plasmids and 50 reference phages. The recall of a dataset was defined as the number of successful runs divided by the number of genomes in the dataset. **(e)** Precision estimates for DomCycle, metaplasmidSPAdes, Recycler and SCAPP on reference plasmids and phages. Precision was defined as the number of successful runs in the dataset divided by the total number of reported cycles in the dataset. DomCycle displays perfect (100%) precision on the reference plasmids and phages tested. **(f)** The number of reported cycles per sample for DomCycle, metaplasmidSPAdes, Recycler and SCAPP when tested on reference plasmids and phages. DomCycle reported at most a single cycle with the tested reference plasmids and phages.

Next, we evaluated the performance of DomCycle on a set of 56 reference plasmids and 50 reference phage genomes (**Supp. Table 2**). The recall was 0.43 for plasmids and 0.86 for phage which was slightly lower than existing tools (**Figure 2d**). Importantly, DomCycle had perfect precision (**Figure 2e**) and consistently reported only a single cycle (or no cycle at all), while the other tools occasionally split a single plasmid or phage into multiple reported elements (**Figure 2f**). To summarize, DomCycle effectively avoided reporting phantom cycles altogether, while achieving recall values that were close to, but slightly inferior than existing tools.

### Recovering dominant cycles with simulated variant mobile elements

To test performance in the face of polymorphisms we simulated two evolutionary scenarios. The first scenario involved a single random plasmid (central allele) and 8 variant plasmids (variant alleles) that were individually distinguished from the major allele by a single random genome rearrangement event (insertion, deletion or inversion). The second scenario, simulating a semi-induced prophage, contained a single prophage that appeared in a circular form (central allele) and integrated 8 times into a single large genome (variant alleles). For both scenarios, we tested recall and precision as a function of central allele frequency. DomCycle successfully recovered 47.3% of central allele plasmids and 39.5% of central allele phage. Central alleles were recovered when the allele frequency surpassed 55% for plasmids and 65% for phage (**Figure 3a**). Despite a background of convoluted genome rearrangements, the precision was perfect in all cases, i.e., all cycles reported by DomCycle were associated with real underlying genomes and multiple cycles were never reported. To illustrate the performance of DomCycle, we show the underlying graph cycle (**Figure 3b**) and the distribution of mapped reads along the cycle (**Figure 3c**) for a single successful plasmid run.

**Figure 3.**
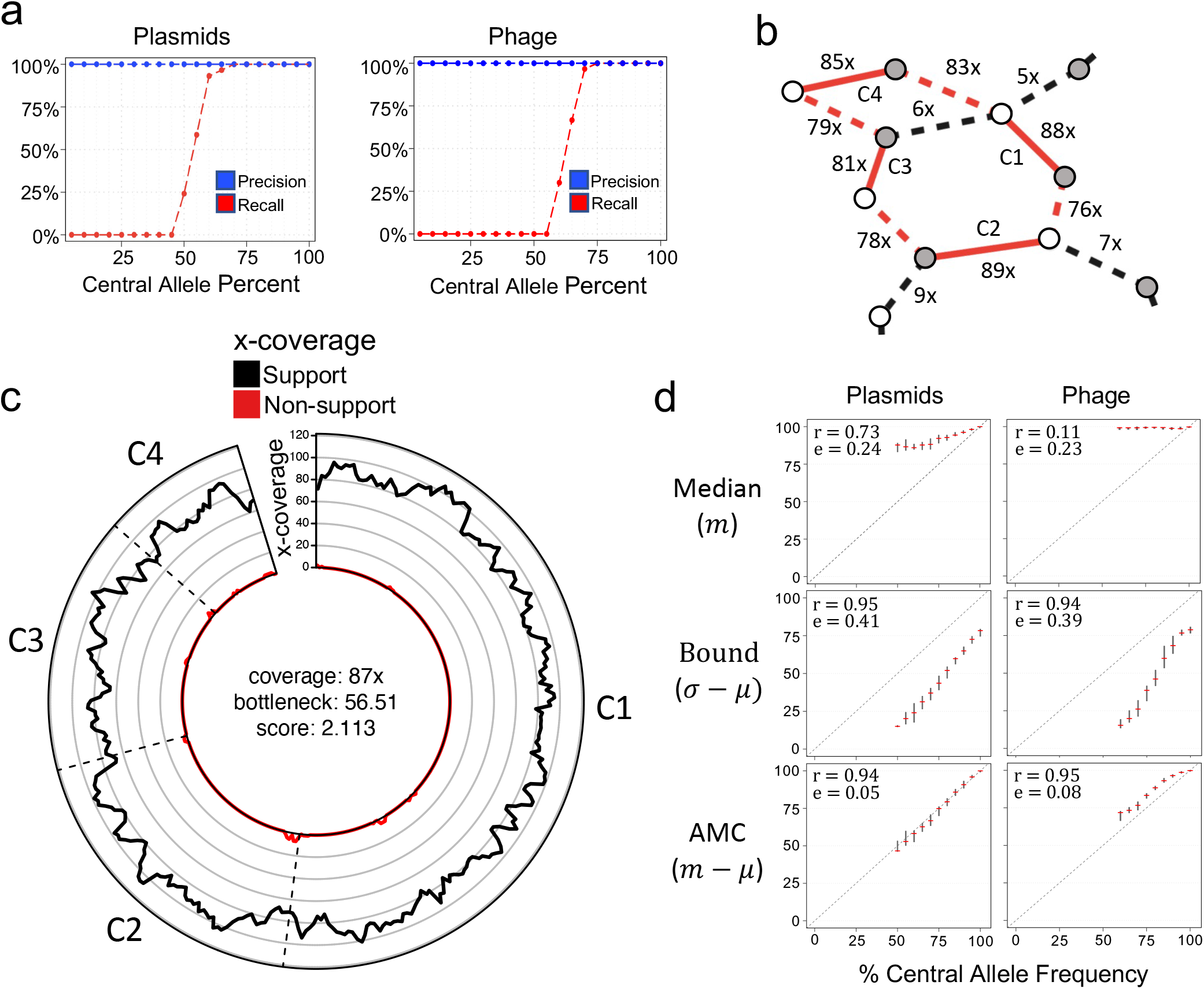
DomCycle performance on recombining plasmids and partially induced phage. **(a)** The recall (red) and precision (blue) for simulations at varying central allele frequencies. Each point represents the results of 30 trials at a single central allele frequency. Central allele frequency was calculated as the percent of the total x-coverage contributed by the central allele. **(b)** Example of a recovered central allele plasmid with a frequency of 55%. White points represent tail vertices and grey points represent head vertices, while internal edges are illustrated with solid lines and external edges are illustrated with dotted lines. The recovered cycle is colored in red and adjacent graph edges that are not part of the cycle are colored in black. Coverage units show the edge coverage, W(e). Labels on internal edges show contig names. **(c)** The nucleotide-level cycle coverage profile corresponding to the cycle depicted in (b). The coverage of cycle supporting reads is colored in black and the coverage of non-supporting reads is colored in red. **(d)** Median coverage, lower bound coverage, and adjusted median coverage (AMC) as predictors of true allele frequency. The Pearson correlation coefficient and root mean squared deviation are shown for each predictor. In each small cross, the red horizontal line shows the median metric value of an estimator at a given central allele frequency, and the vertical line depicts the interquartile range of the estimator, created with 30 replicates for each allele frequency. Diagonal dotted lines show the true central allele frequency.

We leveraged knowledge of the underlying coverages to examine the performance of three genome coverage estimators (**Figure 3d**). The best estimator was the adjusted median coverage (AMC), defined as *x_c_* = *m_c_* − *τ_c_*, where *m_c_* is the median coverage along the cycle. AMC was both highly correlated with true coverage values (Pearson’s coefficient *ρ* = 0.94), and had an RMSD of 0.05 and 0.08 for plasmids and phages respectively. In summary, the analysis of simulated data demonstrated the ability of DomCycle to recover dominant cycles and accurately predict their coverage while faced with complex polymorphic mobile elements.

### Circular genetic elements in the human gut

Next, DomCycle was applied to metagenomic data from stool of a healthy adult (200M paired reads, publicly available data from a previous study^27^). The contig-level and nucleotide-level assessment of cycles was in general agreement (**Supp. Figure S1**), with only 8 cycles that were dropped due to abnormal read coverage that likely stemmed from assembly artifacts (**Supp. Figure S2**). After stringent cycle vetting, 49 dominant cycles and their corresponding ecMGEs remained. Their genome lengths ranged from 0.6kb to 185kb (median length 4.8kb). In terms of complexity, 19% of the ecMGE genomes spanned multiple contigs and 13% of ecMGEs were self-loops that involved bridging an assembly gap using paired reads. We offer an example of a putative phage (**Figure 4a**) and a putative plasmid (**Figure 4b**) (all ecMGEs are shown in **Supp. Figure S3**). Of note, among the recovered ecMGEs was a complete genome of a member of the crAssphage family^27^, a group of prevalent *Bacteroides* phage (**Supp. Figure S4**). The number of cycles identified in this dataset by existing tools was similar to their expected rate of false-positives, suggesting that the low precision of existing tools significantly confounds MGE reporting on real data (**Figure 4c**).

**Figure 4.**
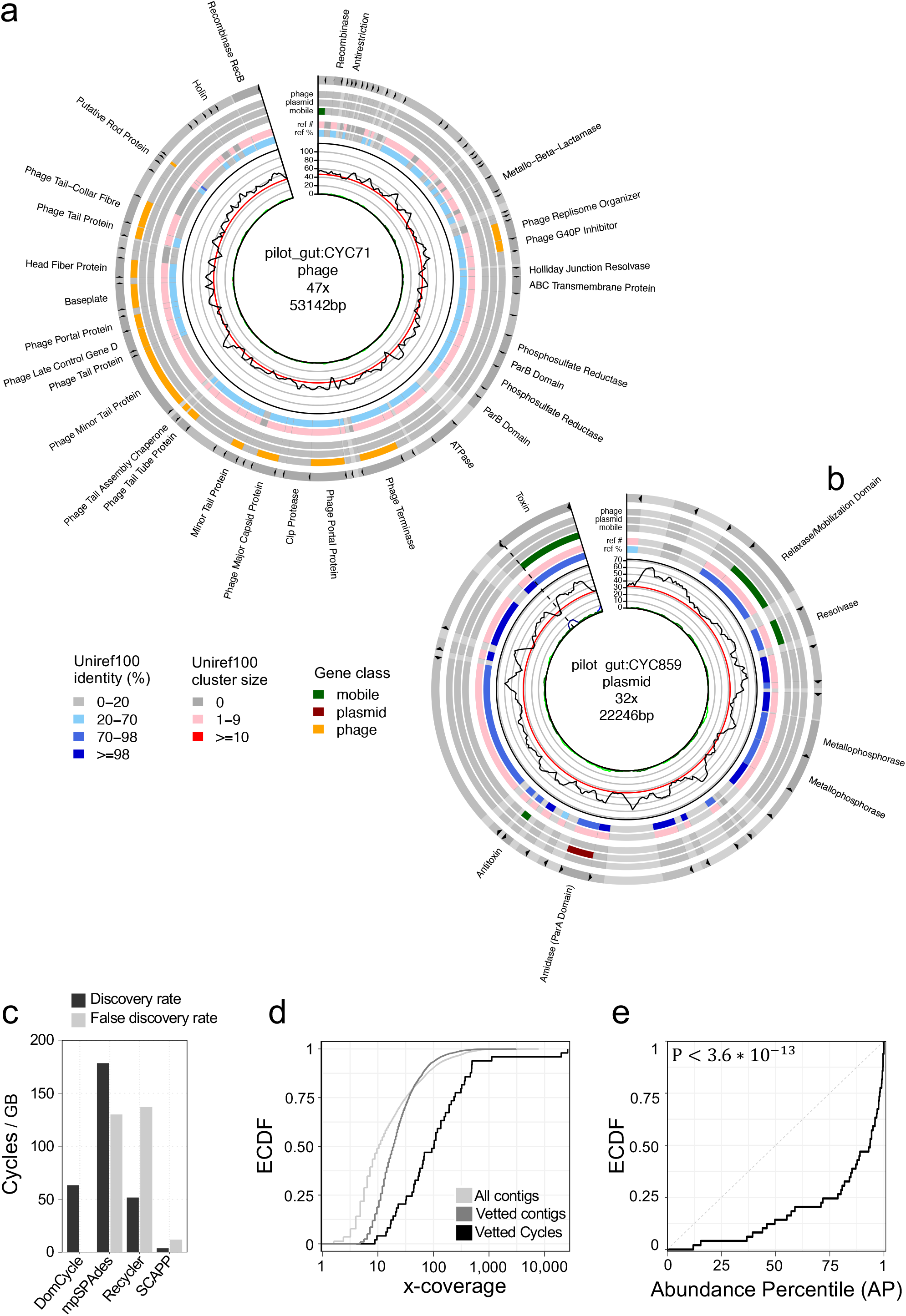
ecMGEs in the gut of a healthy adult. **(a)** A putative phage. Shown from inner to outer circles are nucleotide-level coverage profiles, uniref100 hits (sequence identity (AAI) to the best uniref100 hit and the number of uniport genes in the uniref100 cluster), and gene classification. Gene descriptions for select phage-associated genes are specified outside. Cycles are cut open at the start of their linear sequence for visualization purposes. **(b)** Same as panel a, for a putative plasmid. **(c)** Number of reported cycles per GB of assembly (black) and the estimated rate of false-positives (gray, based on Figure 2b), compared among several tools. **(d)** Empirical cumulative distribution functions (ECDF) for the adjusted median coverage (AMC) of the 49 vetted dominant cycles that correspond to putative ecMGEs (black), the background AMC defined as the AMC of contigs vetted in the same way as dominant cycles except for the circularity condition (dark gray) and the background coverage all contigs in the assembly (light gray). **(e)** Shown for all 49 vetted dominant cycles is the ECDF of the cycle abundance percentile (AP), defined as the percentile of the cycle AMC within the background AMC distribution.

The AMC of the 49 ecMGEs ranged from 8.8x to 25,960x and was on average 10-fold higher than the average coverage of all contigs in the assembly (**Figure 4d**). This suggested that the ecMGEs we detected may have elevated abundance levels compared to the average abundance of bacterial members in the microbial community. However, to properly interpret AMC values we needed to take into account the reduced probability of detecting low-coverage cycles due to our stringent vetting procedure. We computed for each ecMGE its *abundance percentile* (AP), defined as the percentile of its AMC score within a background distribution of AMC values estimated using all contigs in the assembly (**Methods**). Notably, even after this normalization, ecMGEs were significantly abundant within the community (Kolmogorov–Smirnov test *D* = 0.543, *P* < 3.6 ∗ 10^−13^), with 20 ecMGEs (41%) above the 95^th^ abundance percentile (**Figure 4e**). Analysis of a second subject (sequenced with 96M paired reads^28^), in which 20 dominant cycles and associated ecMGEs were recovered, qualitatively recapitulated these results (**Supp. Figure S5**).

### Circular mobile elements are abundant in diverse environments

Next, we applied DomCycle to 30 additional shotgun libraries, including human stool (Gut HMP data^29^, n=10, median 104M reads per sample), sewage wastewater^30^ (n=10, median 48M reads per sample) and the marine environment^31^ (n=10, median 37M reads per sample) (**Supp. Table 3**). In total, we identified 717 dominant cycles and reconstructed their associated ecMGE genomes. Analysis was limited to 221 ecMGEs (29%) that were at least 1kb long (**Figure 5a**). All ecMGEs were classified based on annotations of predicted genes, as putative plasmids (28%), putative phage (16%), unspecified mobile elements (17%) and undefined elements (38%) (**Figure 5b**). The 3 environments differed in the distribution of classes (chi-squared test *P* < 10^−16^), with the gut relatively depleted for undefined elements, the sewage enriched for plasmids and undefined elements, and the ocean enriched for undefined elements (**Figure 5c**). Recapitulating our observation in the two pilot gut samples, the 221 recovered ecMGEs were highly abundant (KS-test *P* < 10^−16^), with 73% of ecMGEs above the 95^th^ abundance percentile within their respective communities. All ecMGE classes were associated to some degree with elevated abundance levels (**Figure 5d**). Elevated abundance of ecMGEs was observed in all environments yet was most prominent in the human gut (**Figure 5e**).

**Figure 5.**
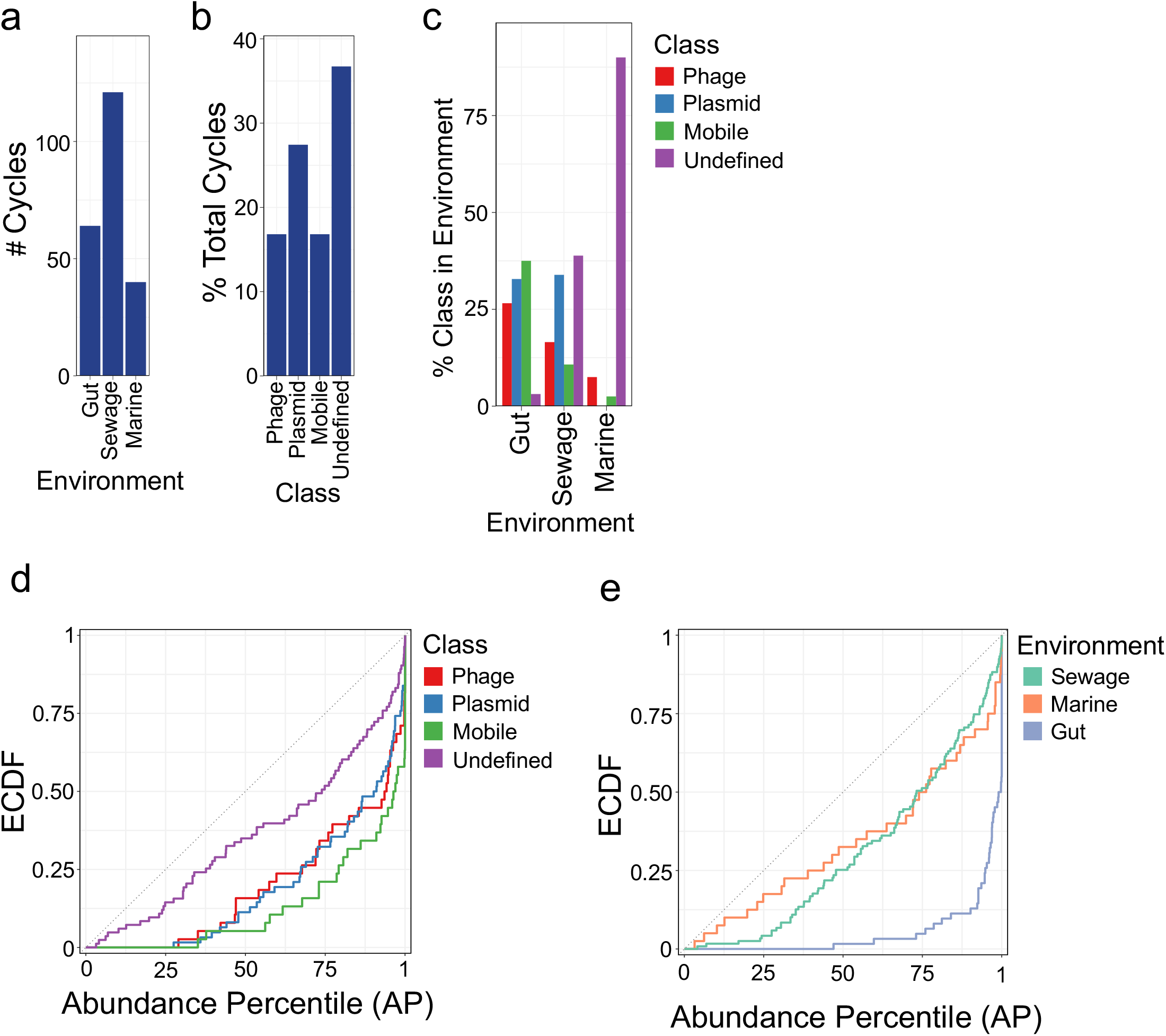
ecMGEs in the human gut, sewage wastewater and marine environments. **(a)** The number of ecMGEs identified in each environment. **(b)** The percentage of ecMGEs with assigned functional classes. **(c)** The percentage of ecMGEs within each environment, stratified by class. **(d)** The ECDF of AP values of ecMGEs, stratified by class. **(e)** Same plot as panel d, stratified by environment.

### Highly prevalent plasmids are rapidly circulating

We were curious to see if we could find evidence of ecMGEs circulating within and between environments. For this analysis, we performed pairwise genome alignments of all 286 ecMGEs (>1kb) detected in the 32 environmental samples described in this work. A comparison of the fraction of the aligned region and the average nucleotide identity (ANI) within the aligned region suggested that their genome structure is highly conserved (**Figure 6a**). The ecMGEs were grouped into 244 clusters based on sequence identity using a threshold of 95% ANI (**Supp. Figure S6**). Results were robust to changes in the clustering threshold (**Supp. Figure S7**). Analysis was limited to the 20 clusters (denoted M1-M20) that had two or more members (**Table 1**; See **Supp. Table 4** for cluster members). Clusters were extremely tight; the average fraction of the aligned region between pairs of cluster members was 99.56-100% (median 100%) and the average ANI within the aligned region was 98.43-100% (median 99.9%). We refer to these clustered ecMGEs, which were reconstructed independently with minor genetic variations in multiple samples, as *circulating* elements.

**Figure 6.**
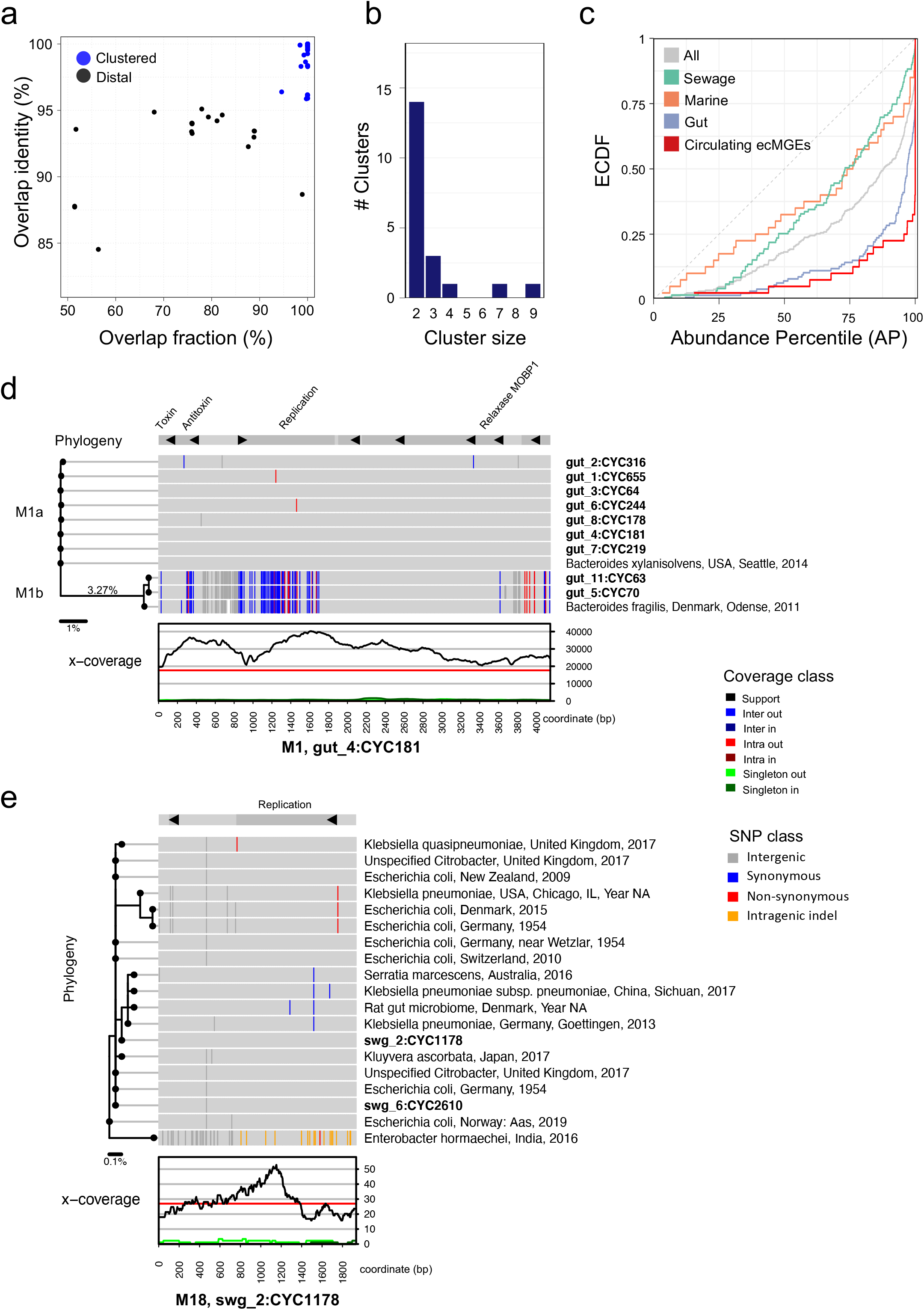
Analysis of circulating ecMGEs. **(a)** The distribution of cycle tightness metrics across pairs of ecMGEs (>1kb) in the 32 samples assayed. The overlap identity is defined as the average identity of aligned regions between two cycle genomes. The overlap fraction is defined as the percent of two cycle genomes that align. For a cycle pair, genomic similarity is defined as the overlap identity times the overlap fraction. Cycle pairs are linked if their genomic similarity is >0.95 and clusters are defined as groups of linked cycles. **(b)** The distribution of the number of members in clusters. **(c)** The ECDF of AP values of circulating ecMGEs, compared to ecMGEs stratified by their environment and to all ecMGEs. **(d)** Detailed view of cluster M1. Data are projected onto a reference ‘pivot’ cluster member (gut4, cycle 181) that is shown in a linear format for visualization purposes. Shown from the bottom up are the x-coverage profiles of the pivot ecMGE member (see color legend), SNP patterns of cluster members (in bold) and reference sequences (isolate source, location and year of collection), and annotated genes on top. SNP patterns are colored according to differences from the pivot, with white indicating segments that failed to align. A phylogenic tree is shown on the left. The units of the scale bar under the tree are mean nucleotide differences. Clades M1a and M1b are marked on the plot. **(e)** Detailed view of cluster M18, see panel (d) for a description.

**Table 1.** Table columns from left to right specify for each cluster the number of cluster members, the average genome length in base pairs, the environments in which the cluster was identified (out of the 32 samples examined), the average nucleotide identity (ANI) between pairs of cluster members, the average abundance percentile, the cluster classification, number of genes, number of genes that did not match the queried databases or matched an uncharacterized or hypothetical protein, mobility classification based on HMMs, presence of replication gene, presence of a toxin gene (T) and/or an anti-toxin gene (A), number of hits (>95% ANI) in the plasmid database PLSDB, and the taxonomic families as inferred from the isolate source specified in PLSDB.

| id | #members | length | environment | identity | AP | class | #genes | #uncharacterized | mobility | replication | addiction | #references | host family |
| --- | --- | --- | --- | --- | --- | --- | --- | --- | --- | --- | --- | --- | --- |
| M1 | 9 | 4148 | Gut | 98.27 | 0.999 | plasmid | 8 | 4 | MOBP1 | Y | T+A | 2 | Bacteroidaceae |
| M2 | 7 | 2751 | Gut | 98.97 | 0.998 | plasmid | 2 | 1 | MOBP1 |  |  | 3 | Bacteroidaceae<br>Enterobacteriaceae |
| M3 | 4 | 4494 | Gut, Sewage | 99.92 | 0.939 | plasmid | 8 | 4 | NA | Y | T+A | 6 | Bacteroidaceae<br>Moraxellaceae |
| M4 | 3 | 5820 | Gut | 99.94 | 1 | plasmid | 9 | 5 | MOBP1 | Y | T+A | 0 |  |
| M5 | 3 | 4971 | Gut | 99.7 | 0.947 | plasmid | 8 | 2 | MOBP1 | Y | T | 0 |  |
| M6 | 3 | 2413 | Gut | 98.86 | 0.366 | undetermined | 4 | 3 | NA |  |  | 0 |  |
| M7 | 2 | 3034 | Gut | 99.9 | 0.7076 | plasmid | 2 | 0 | MOBV | Y |  | 0 |  |
| M8 | 2 | 2775 | Gut | 99.61 | 0.804 | plasmid | 2 | 0 | MOBV | Y |  | 1 |  |
| M9 | 2 | 2750 | Gut | 98.4 | 1 | plasmid | 2 | 0 | MOBP1 | Y |  | 3 | Pseudomonadaceae<br>Bacteroidaceae |
| M10 | 2 | 8598 | Gut | 99.52 | 0.837 | plasmid | 11 | 5 | MOBF |  |  | 0 |  |
| M11 | 2 | 5843 | Gut | 99.87 | 0.389 | plasmid | 7 | 4 | MOBP1 |  |  | 0 |  |
| M12 | 2 | 1488 | Sewage | 100 | 0.408 | plasmid | 2 | 1 | NA | Y |  | 2 | Enterobacteriaceae |
| M13 | 2 | 1483 | Sewage | 99.26 | 0.551 | plasmid | 2 | 1 | NA | Y |  | 0 |  |
| M14 | 2 | 3086 | Sewage | 100 | 0.546 | plasmid | 3 | 1 | MOBQ |  |  | 0 |  |
| M15 | 2 | 1044 | Marine | 99.61 | 0.899 | undetermined | 0 | 0 | NA |  |  | 0 |  |
| M16 | 2 | 2585 | Sewage | 100 | 0.947 | plasmid | 3 | 0 | MOBV |  | A | 0 |  |
| M17 | 2 | 2349 | Sewage | 99.91 | 0.536 | plasmid | 2 | 0 | MOBP1 | Y |  | 0 |  |
| M18 | 2 | 1934 | Sewage | 99.95 | 0.389 | plasmid | 2 | 1 | NA | Y |  | 17 | Enterobacteriaceae<br>Yersiniaceae |
| M19 | 2 | 1382 | Sewage | 100 | 0.613 | plasmid | 2 | 1 | MOBT | Y |  | 0 |  |
| M20 | 2 | 1705 | Sewage | 98.22 | 0.482 | plasmid | 1 | 0 | NA | Y |  | 0 |  |

We classified 16 clusters as putative plasmids due to the presence of mobility and/or replication genes, of which 5 had one or two toxin-antitoxin genes, as summarized in Table 1 (details in **Supp. Table 5**). There were 33 uncharacterized genes in total, which made up 32% of each cluster on average (**Supp. Table 6**). Clusters had 2-9 ecMGE members (**Figure 6b**), and all were from a single environment except cluster M3 that was observed in both gut and sewage samples. The abundance percentile (AP) of circulating ecMGEs was high (mean AP of 92%), in agreement with a commonly observed ecological association between prevalence and abundance (**Figure 6c**).

Leveraging a comprehensive plasmid reference database (PLSDB^32^, with over 26k plasmids), we identified 7 clusters with reference hits (99.7% ANI on average, **Supp. Table 7**). This independent (reference) reconstruction of the circulating ecMGEs provided validation of our metagenomic approach. Moreover, the isolate source information informed us about the putative host-range and environments of some of our clusters. These data suggested that our circulating ecMGEs may have a broad host-range, with 5 (out of 6 with host data) associated with multiple taxonomic families (Table 1). We discovered that the top three plasmids (M1-M3) were reported as cryptic *Bacteroides* plasmids in the 1980s^33,34^. The 13 circulating ecMGEs that did not have a close reference in PLSDB (>95% ANI) despite their prevalence in the general population highlight the discovery power of our metagenomic approach.

We proceeded to augment clusters with reference genomes (where available) and inferred cluster-specific phylogenic trees. The most prevalent plasmid (M1, 4138+/−10bp) was observed in 9 gut samples and had 2 isolate-based reference genomes (**Figure 6d**). M1 is composed of a major clade (M1a, isolated previously from *B. xylanisolvens*) and a minor clade (M1b, isolated previously from *B. fragilis*), with a genetic distance of 3.27% separating the two inferred clade ancestors. Both have shallow clonal trees distinguished by a handful of SNPs, with a mean ANI of 99.95% and 99.76% between clade members for M1a and M1b respectively. Out of the 12 gut samples we assayed, M1a was present in 7 out of the 12 gut samples (58+/−22%) making it one of the most prevalent plasmids recorded in the human gut to date. The most recent common ancestor is also relatively recent on an evolutionary timescale; a coalescence analysis suggests M1a may have gone through a clonal expansion merely ~600-1200 years ago (**Supp. Note 4**).

Another cluster of interest is M18, a short cryptic plasmid (1934bp) recovered from 2 sewage metagenomic samples and independently reconstructed from 22 isolates (primarily *Enterobacteriaceae* species) that were collected across the globe (**Figure 6e**). The clonal population structure of M18 is in discordance with its host species, suggesting it is freely circulating between a diverse set of hosts. The remaining 5 ecMGE clusters that have reference genomes suggest that ecMGEs are found in diverse environments and bacterial hosts (**Supp. Figure S8)**. The 9 ecMGE clusters with sufficient members to estimate their phylogeny had clonal populations with only 0-3.2% putative recombined sites (**Supp. Figure S9**). Finally, we noted several examples of clusters with an uneven distribution of SNPs along their genomes, suggestive of possible adaptive evolution or recombination (**Supp. Figure S10**).

## DISCUSSION

In this work, we present an algorithm that recovers all dominant cycles in a metagenomic assembly graph and reconstructs their corresponding genomes. Our implementation achieves high precision by combining graph theory and nucleotide-level vetting of cycles. We show that in the context of microbial communities, dominant cycles likely correspond to true extrachromosomal circular DNA. Application to complex evolutionary scenarios and reference data reliably recovers ecMGEs without reporting any false-positives.

The approach recovers only circular genomes that correspond to graph cycles. In reality, circular mobile elements can contain long repeat elements (>77bp, which is the kmer size used during the assembly) and will therefore produce complex graph circuits that would not be detected. (Unlike a cycle, the path of a circuit can traverse the same contig more than once.) The ongoing transition to long-read sequencing technologies is expected to alleviate this problem by transforming some complex circuits to cycles. With meticulous handling of environmental samples, long-reads can extend up to 5-10kb^25,35^. Combining long-reads with the approach presented here will allow genotyping of complex ecMGEs, such as MGEs that contain insertion sequences (typically <2.5kb) and short transposons. Longer repeat elements and complex rearrangements that require bridging over more than ~10kb can be addressed with additional experimental work, such as Hi-C^28,36–38^ and by sampling the same community multiple times^39,40^.

Application of our approach to 32 environmental samples uncovered 20 clades of ecMGEs (primally gut and sewage plasmids), showcasing the strength of metagenomic approaches in tapping into understudied environmental plasmids. The surprisingly low sequence diversity and clonal population structure we report here were recently observed in plant-associated virulence plasmids^41^. Plasmid clonality is particularly striking when contrasted with the population structure of their bacterial hosts, which partake in pervasive recombination at levels that can obscure strain phylogeny^42–44^. The clonality and lack of diversity can be partially attributed to their simplified ecological niche that may favor rapid cycles of selective sweeps. The relative conservation of their genome structure (as shown in Figure 6a) suggests that among their uncharacterized genes, some might support the prolific lifestyle of these circulating plasmids. Further characterization of the adaptive landscape of MGEs in the gut and elsewhere will require a larger dataset. In summary, this work presents a new tool that allows reconstruction of ecMGEs from readily available public metagenomic shotgun data and that may help to elucidate the evolution and dissemination of mobile genetic elements within and between environments.

## ACKNOWLEDGMENTS

We wish to thank Les Dethlefsen and Benjamin Good for their insightful comments. Research reported in this publication was supported by the National Institute of Allergy and Infectious Diseases of the National Institutes of Health (grant number R01AI147023 to D.A.R.) and the Thomas C. and Joan M. Merigan Endowment at Stanford University (to D.A.R.).

---

**Supplementary Figure 1.**
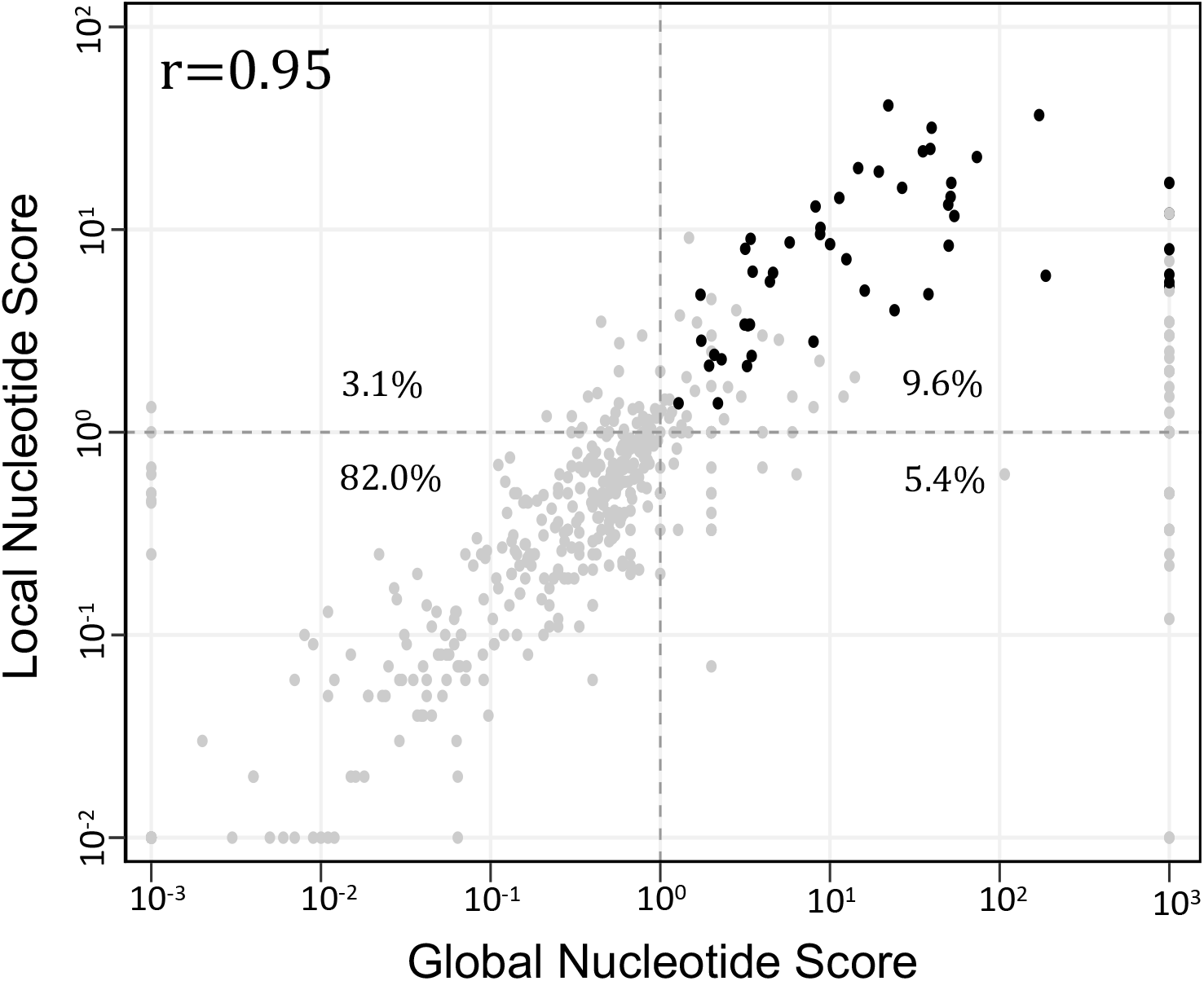
The distribution of the global nucleotide score and local cycle score for each candidate cycle reported on the gut sample from a healthy adult (SRR8187104). Vetted dominant cycles are shown in black and candidate cycles filtered out by either the global nucleotide score test or the local cycle score test are shown in grey. Horizontal dotted lines drawn show the lower-bound threshold for classifying vetted dominant cycles without accounting for significance through p-values in both score tests. The Pearson correlation coefficient is computed between candidate cycle’s global nucleotide score and local cycle score.

**Supplementary Figure 2.**
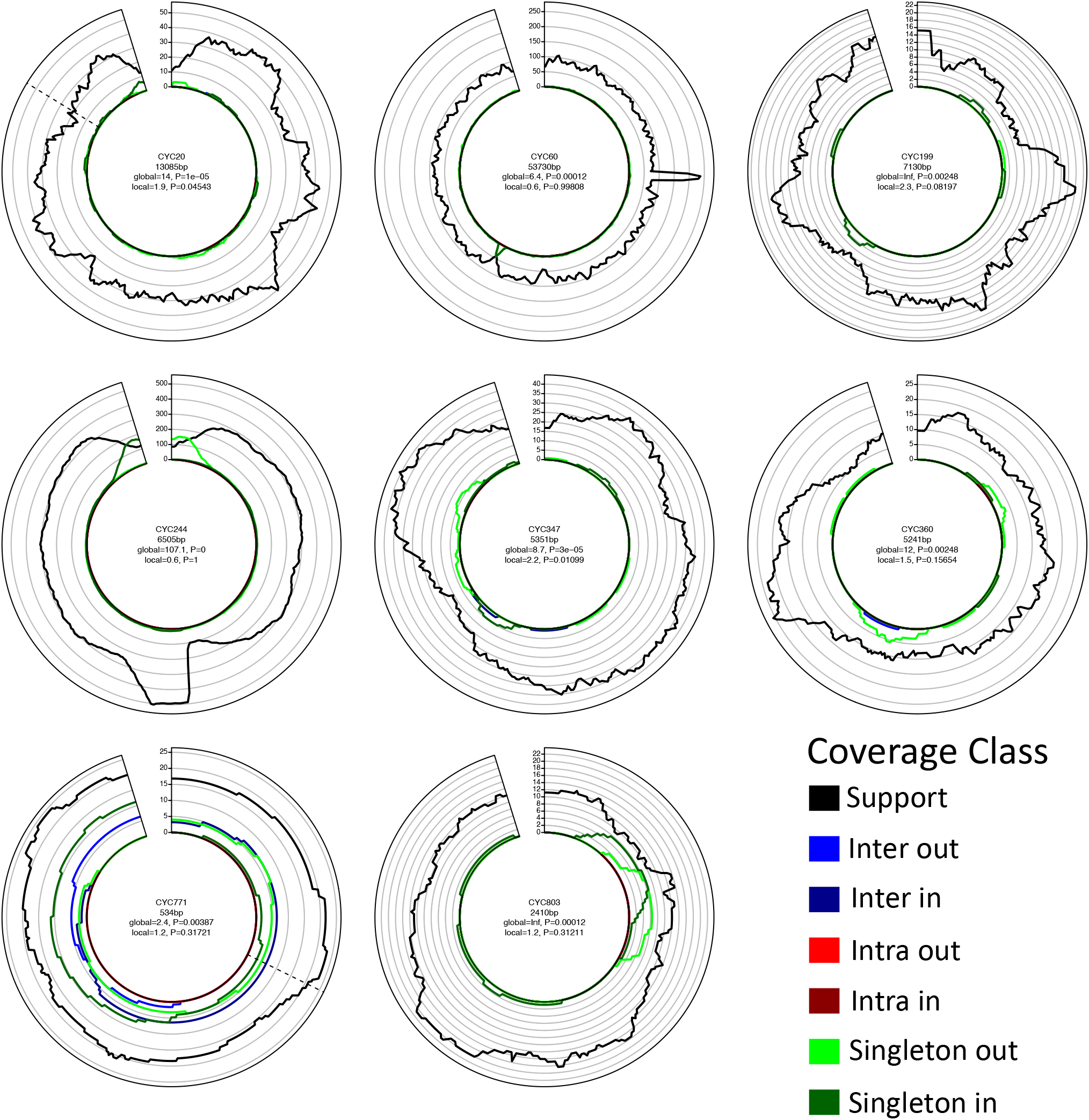
Examples of candidate cycles filtered out due to poor local nucleotide-level scores in the gut of a healthy adult (SRR8187104). The local nucleotide-level score test calculates the p-value for the hypothesis that the support coverage is greater than the total base pair non-support profile at each base in the candidate cycle (see **Methods**). In comparison to the global nucleotide-level score, the local score accounts for the singleton coverage and tests significance at each candidate cycle base. For instance, CYC244 (middle left) has out singleton coverage (light green) that exceeds the support coverage at the beginning of the cycle; accordingly, this cycle receives a non-significant local nucleotide-level score (*p* > 0.01) at the bases where the singleton out coverage exceeds the support. Intuitively, the high density of singleton reads on CYC244 indicates that there was assembly fragmentation near the contig junction at the beginning (and end) of the cycle. We conservatively assume that the missing singleton read side originated from a sequence that is missing in the assembly. Thus, we cannot confidently conclude that CYC244 is dominant. A similar rationale extends to other candidate cycles depicted.

**Supplementary Figure 3.**
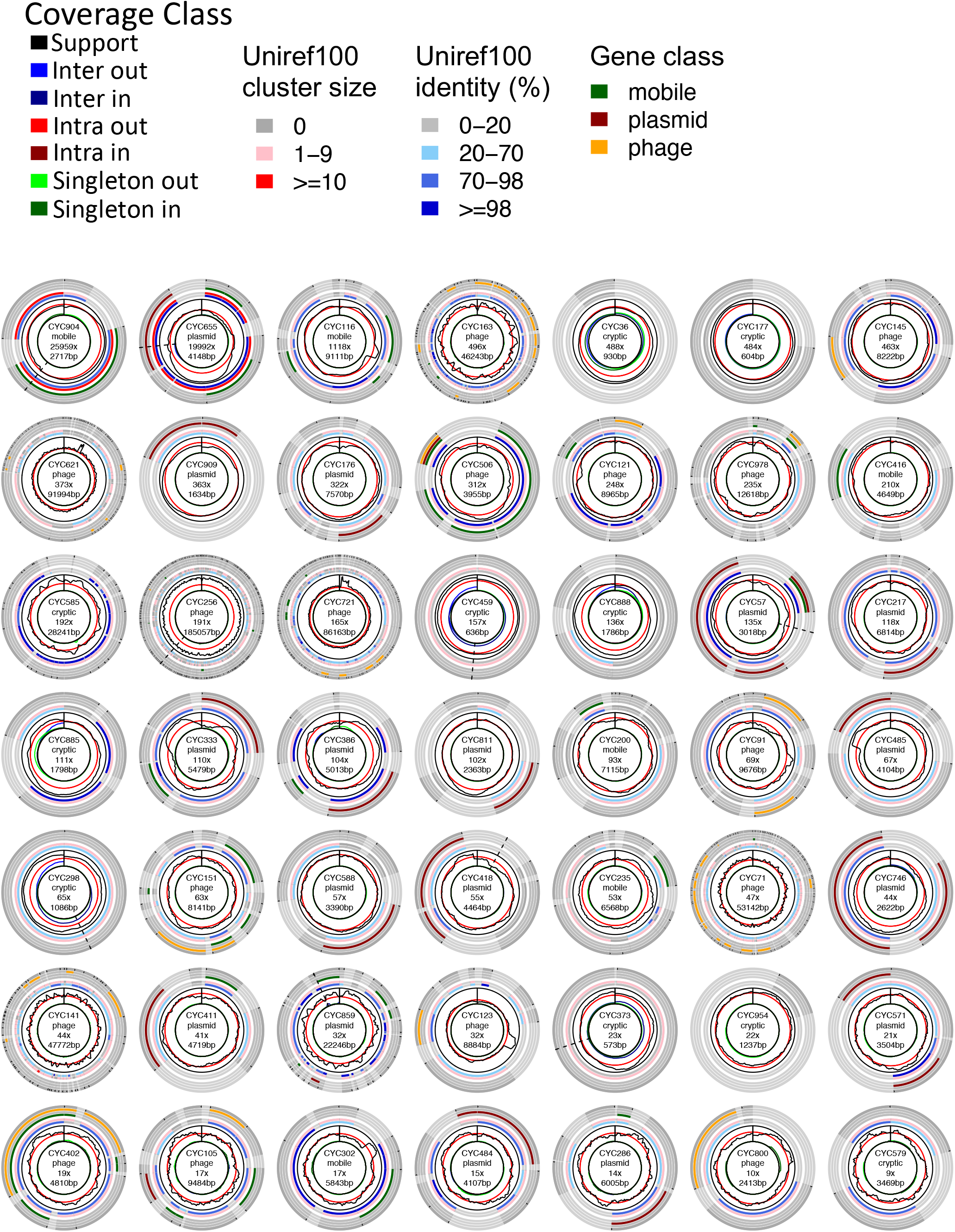
All ecMGEs identified in the gut of pilot subject 1. Each plot shows the coverage profile, gene positions, gene identity, Uniref cluster size, and gene classification for each cycle.

**Supplementary Figure 4.**
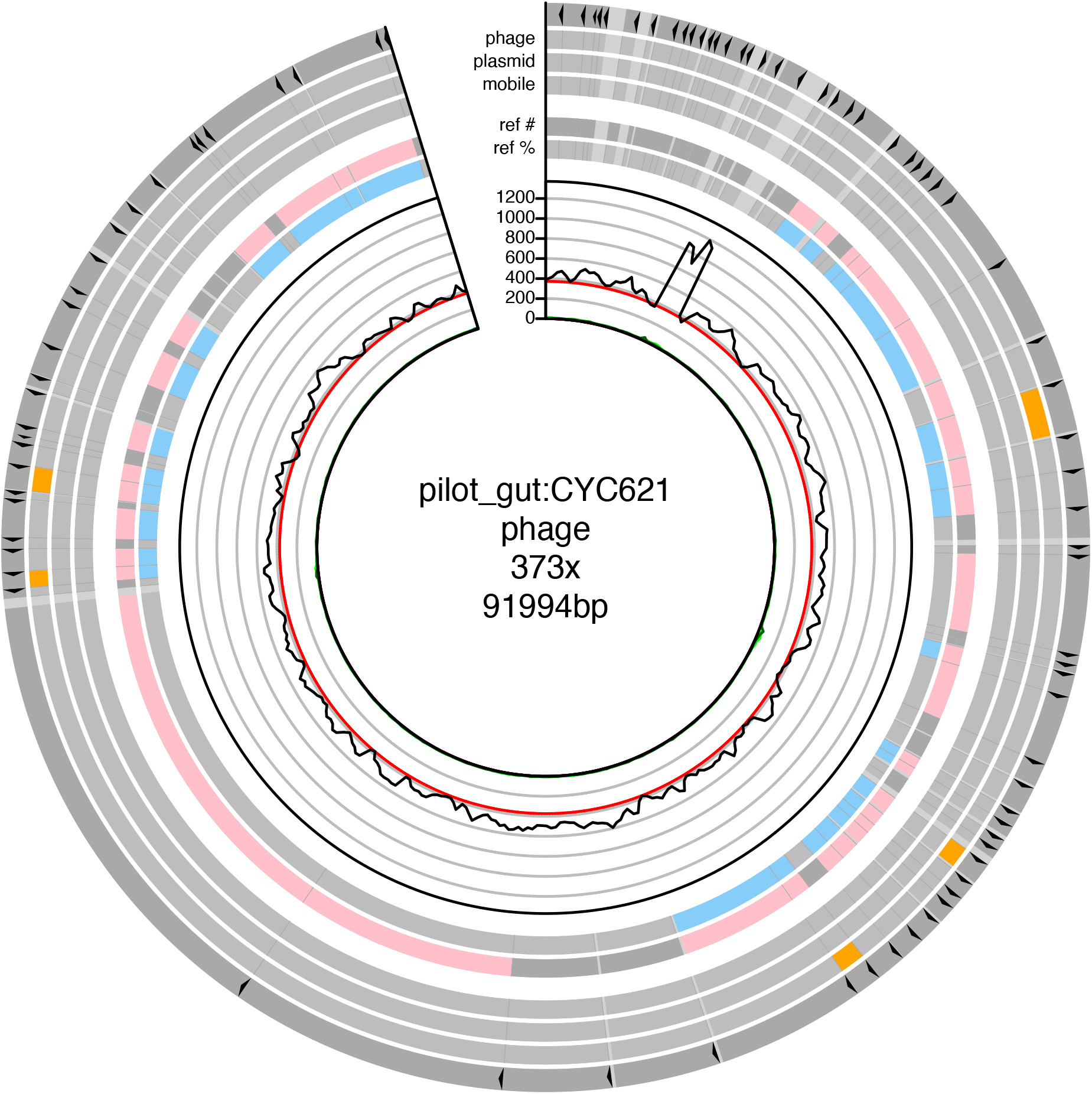
CrAssphage-like element identified in the gut of the central subject in the study.

**Supplementary Figure 5.**
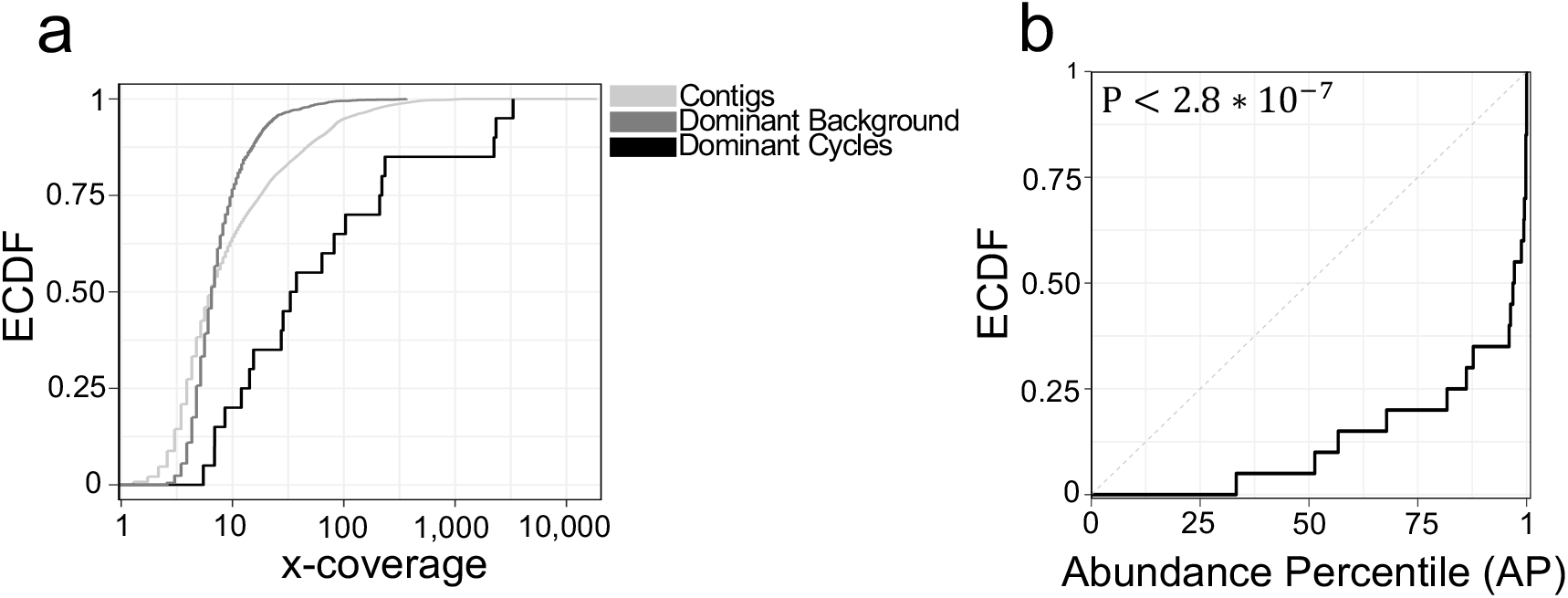
Results for a second deeply-sequenced gut sample from a healthy human adult (SRR8186375) recapitulate trends shown in Figure 4. **(a)** Empirical cumulative density functions (ECDF) for the median support coverage distribution for candidate pseudodominant genomes (light grey), the adjusted median coverage (AMC) distribution for pseudodominant genomes (dark grey), and AMC distribution for dominant cycles (black). **(b)** ECDF for the abundance percentiles (AP) of dominant cycles. The AP for each cycle is computed to be the percentile of the cycle AMC in the background distribution of AMCs among pseudo-dominant genomes. The dotted line shows the AP ECDF for pseudo-dominant genomes (KS-test, P < 2.8 ∗ 10^−7^).

**Supplementary Figure 6.**
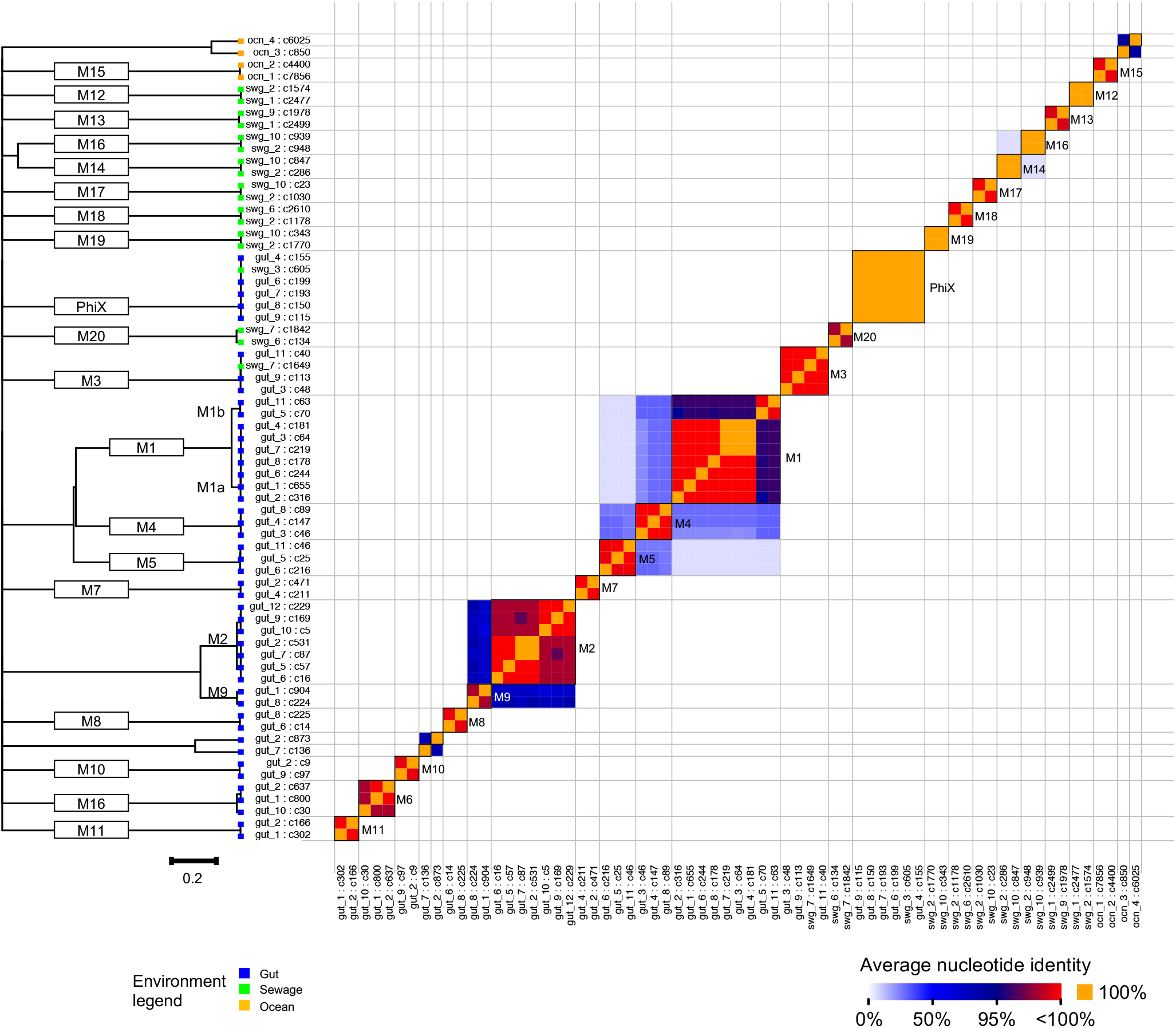
The 286 eMGEs were clustered using hierarchical clustering performed with single linkage. Shown are eMGEs for which the distance to their nearest neighbor was under 0.5 (i.e., >50% ANI). Left shows clustering dendrogram, with a scale bar showing a distance of 0.2 (equivalent to 80% ANI) and nodes colored by environment. Matrix squares colored by sequence identity, with perfect alignments (100% ANI) highlighted in orange. The 20 multi-member clusters (threshold 95% ANI), numbered M1 to M20, are marked on the plot. The PhiX cluster and all single-member clusters were omitted from downstream analysis.

**Supplementary Figure 7.**
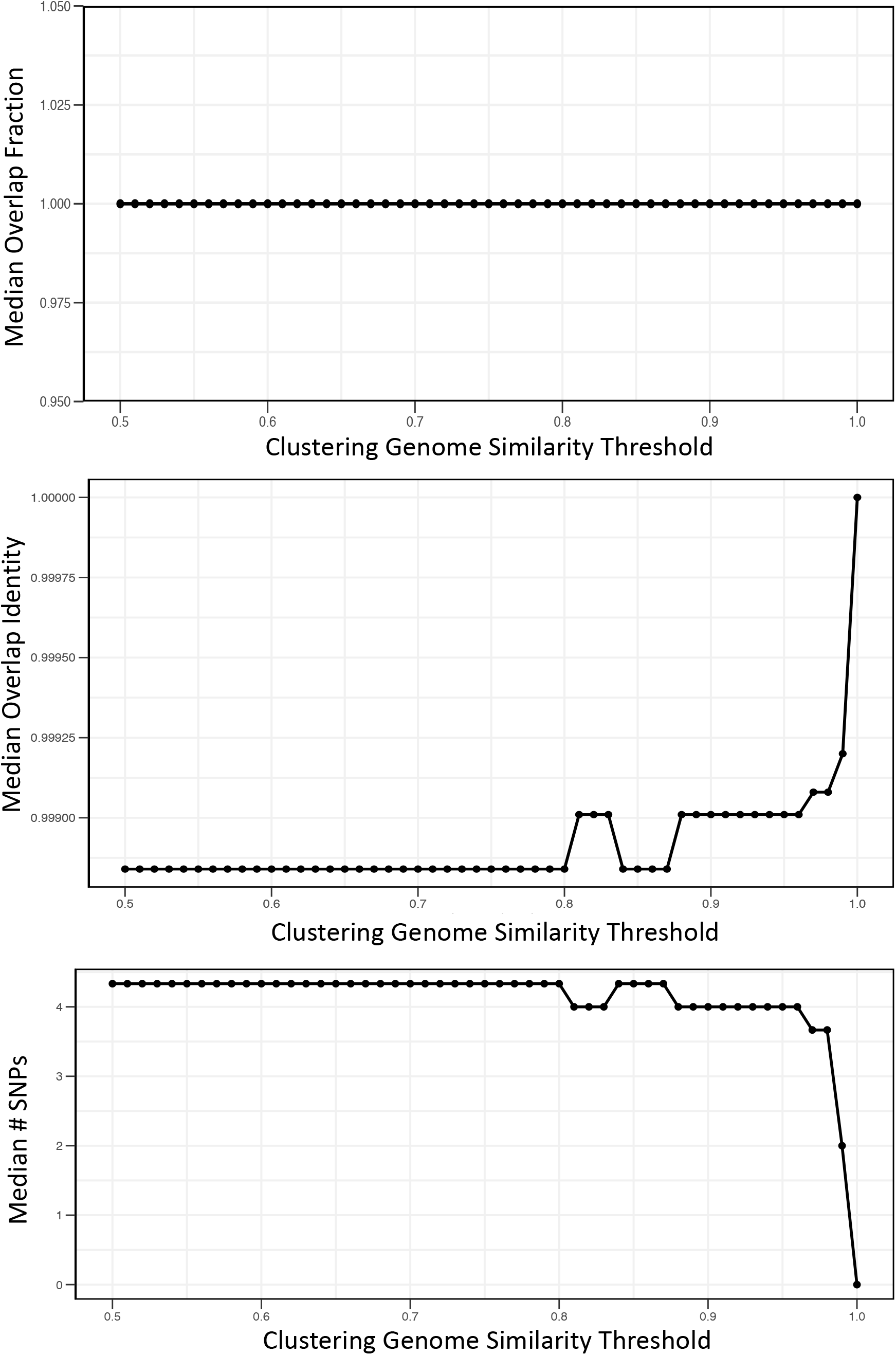
The median of each intra-cluster mean metric as a function of clustering genomic similarity threshold. The three metrics for cluster tightness are robust to changes in the minimum genome similarity clustering threshold.

**Supplementary Figure 8.**
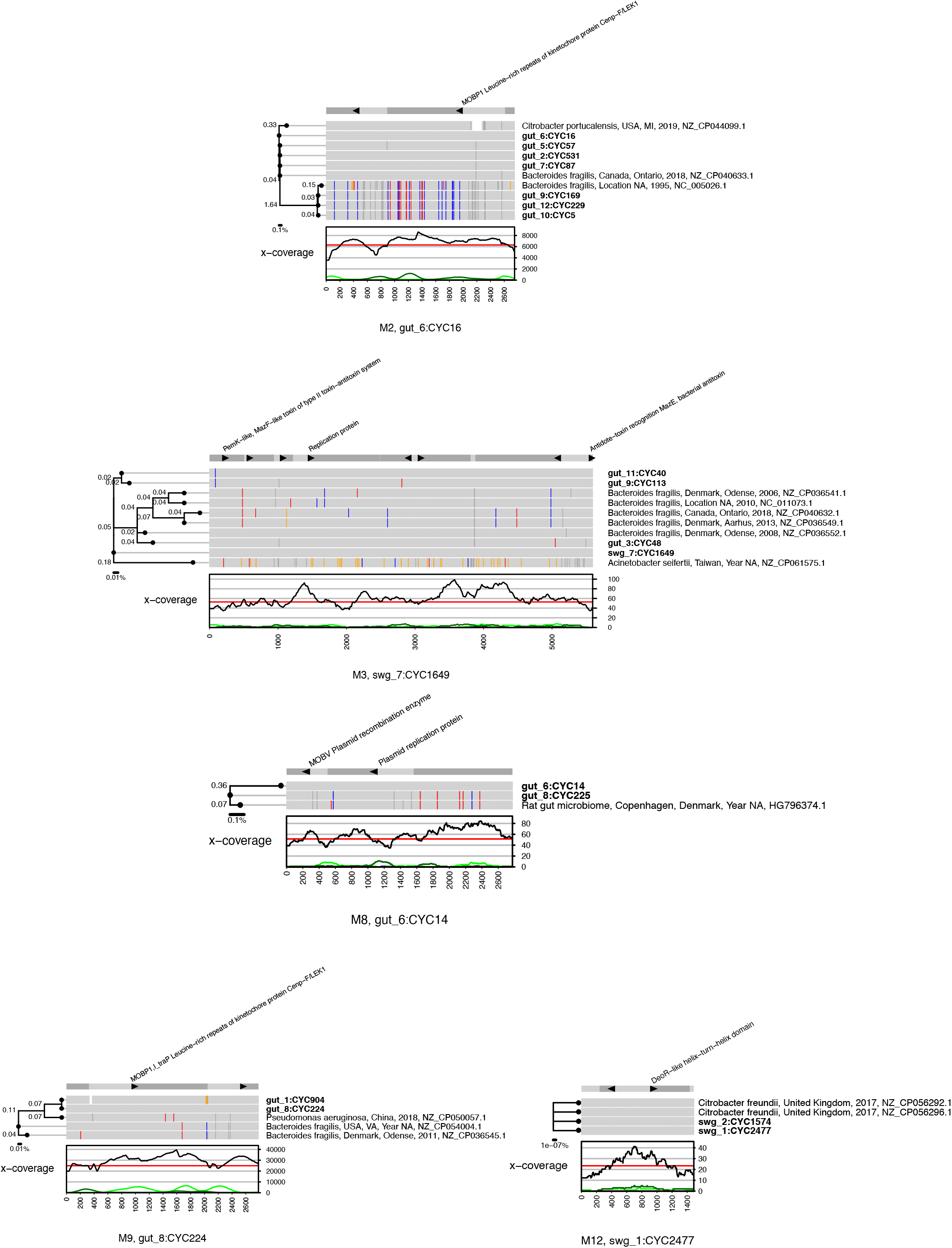
Clusters with one or more references in PLSDB. Legends as in Figure 6. The 5 clusters were found in multiple environments (human gut, sewage and rat microbiome) and are associated with diverse microbial hosts.

**Supplementary Figure 9.**
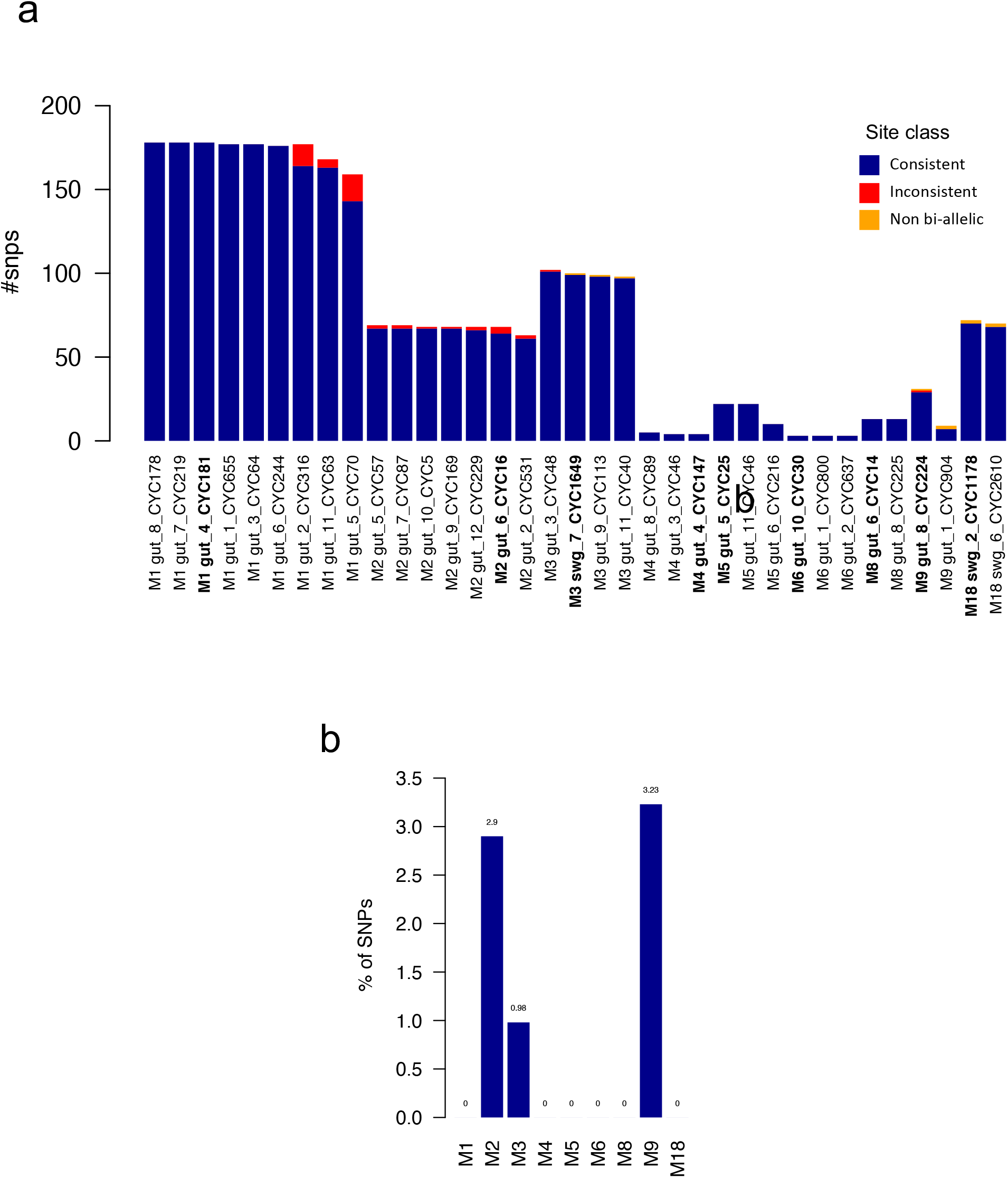
Putative recombination events are rare. For each cluster and each cluster member, the phylogenetic tree as inferred by PhyML was used to classify all polymorphic sites. A bi-allelic site was classified as *consistent* if the partitioning of samples matched an edge in the tree, as *inconsistent* otherwise. Non bi-allelic sites were classified as such. **a)** The breakdown of site classification for all clusters and all options of pivot cluster members. The pivot members selected for visualization purposes in Figure 6 and Supp. Figure 8 and 10 are highlighted in bold. **b)** The maximal percentage of consistent sites out of all sites over all cluster members. Clusters with a value of zero (such as M1) have at least one tree topology that is consistent with all polymorphic sites.

**Supplementary Figure 10.**
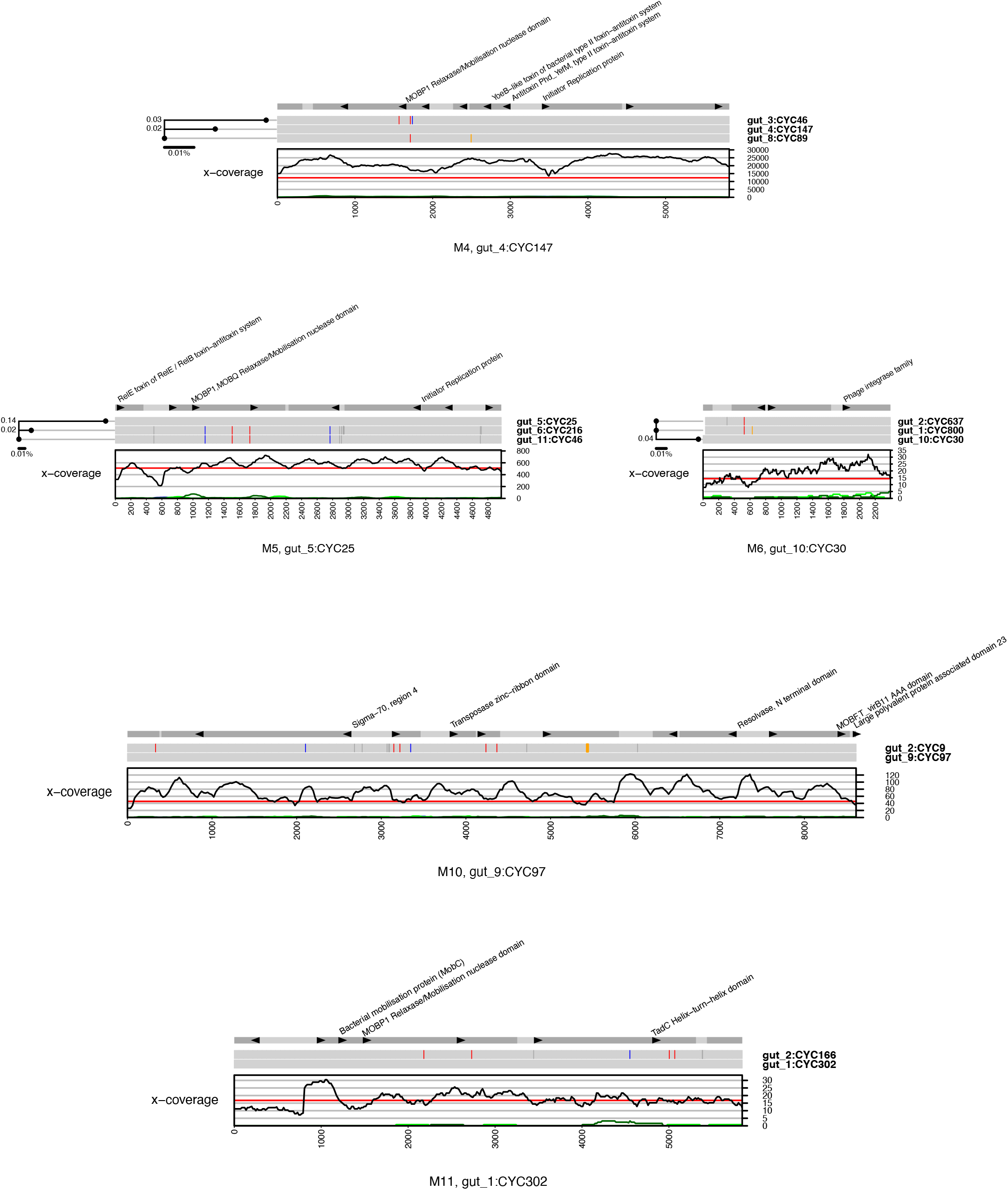
Gut clusters with uneven distribution of SNPs along genomes. Groups of nearby SNPs can indicate a recombination event or positive selection. Legends as in Figure 6.

## REFERENCES

1. Soucy, S. M., Huang, J. & Gogarten, J. P. Horizontal gene transfer: building the web of life. Nat Rev Genet 16, 472–482 (2015).

2. Wiedenbeck, J. & Cohan, F. M. Origins of bacterial diversity through horizontal genetic transfer and adaptation to new ecological niches. FEMS Microbiology Reviews 35, 957–976 (2011).

3. Polz, M. F., Alm, E. J. & Hanage, W. P. Horizontal gene transfer and the evolution of bacterial and archaeal population structure. Trends Genet 29, 170–175 (2013).

4. Deng, Y. et al. Horizontal gene transfer contributes to virulence and antibiotic resistance of Vibrio harveyi 345 based on complete genome sequence analysis. BMC Genomics 20, 761 (2019).

5. Maiques, E. et al. beta-lactam antibiotics induce the SOS response and horizontal transfer of virulence factors in Staphylococcus aureus. J Bacteriol 188, 2726–2729 (2006).

6. von Wintersdorff, C. J. H. et al. Dissemination of Antimicrobial Resistance in Microbial Ecosystems through Horizontal Gene Transfer. Front Microbiol 7, 173 (2016).

7. Frost, L. S., Leplae, R., Summers, A. O. & Toussaint, A. Mobile genetic elements: the agents of open source evolution. Nat Rev Microbiol 3, 722–732 (2005).

8. Sentchilo, V. et al. Community-wide plasmid gene mobilization and selection. ISME J 7, 1173–1186 (2013).

9. Shkoporov, A. N. et al. Reproducible protocols for metagenomic analysis of human faecal phageomes. Microbiome 6, 68 (2018).

10. Jørgensen, T. S., Xu, Z., Hansen, M. A., Sørensen, S. J. & Hansen, L. H. Hundreds of circular novel plasmids and DNA elements identified in a rat cecum metamobilome. PLoS One 9, e87924 (2014).

11. Zhou, F. & Xu, Y. cBar: a computer program to distinguish plasmid-derived from chromosome-derived sequence fragments in metagenomics data. Bioinformatics 26, 2051–2052 (2010).

12. Carattoli, A. et al. In Silico Detection and Typing of Plasmids using PlasmidFinder and Plasmid Multilocus Sequence Typing. Antimicrob. Agents Chemother. 58, 3895–3903 (2014).

13. Roux, S., Enault, F., Hurwitz, B. L. & Sullivan, M. B. VirSorter: mining viral signal from microbial genomic data. PeerJ 3, e985 (2015).

14. Roosaare, M., Puustusmaa, M., Möls, M., Vaher, M. & Remm, M. PlasmidSeeker: identification of known plasmids from bacterial whole genome sequencing reads. PeerJ 6, e4588 (2018).

15. Robertson, J. & Nash, J. H. E. MOB-suite: software tools for clustering, reconstruction and typing of plasmids from draft assemblies. Microbial Genomics 4, (2018).

16. Lanza, V. F. et al. Plasmid flux in Escherichia coli ST131 sublineages, analyzed by plasmid constellation network (PLACNET), a new method for plasmid reconstruction from whole genome sequences. PLoS Genet 10, e1004766 (2014).

17. Conlan, S. et al. Single-molecule sequencing to track plasmid diversity of hospital-associated carbapenemase-producing Enterobacteriaceae. Sci Transl Med 6, 254ra126 (2014).

18. Antipov, D. et al. plasmidSPAdes: assembling plasmids from whole genome sequencing data. Bioinformatics 32, 3380–3387 (2016).

19. Antipov, D., Raiko, M., Lapidus, A. & Pevzner, P. A. Plasmid detection and assembly in genomic and metagenomic data sets. Genome Res 29, 961–968 (2019).

20. Rozov, R. et al. Recycler: an algorithm for detecting plasmids from de novo assembly graphs. Bioinformatics 33, 475–482 (2017).

21. Arredondo-Alonso, S., Willems, R. J., van Schaik, W. & Schürch, A. C. On the (im)possibility of reconstructing plasmids from whole-genome short-read sequencing data. Microb Genom 3, e000128 (2017).

22. Hülter, N. et al. An evolutionary perspective on plasmid lifestyle modes. Current Opinion in Microbiology 38, 74–80 (2017).

23. Kupczok, A. et al. Rates of Mutation and Recombination in Siphoviridae Phage Genome Evolution over Three Decades. Mol Biol Evol 35, 1147–1159 (2018).

24. He, S. et al. Insertion Sequence IS26 Reorganizes Plasmids in Clinically Isolated Multidrug-Resistant Bacteria by Replicative Transposition. mBio 6, e00762 (2015).

25. Suzuki, Y. et al. Long-read metagenomic exploration of extrachromosomal mobile genetic elements in the human gut. Microbiome 7, 119 (2019).

26. Pellow, D. et al. SCAPP: An algorithm for improved plasmid assembly in metagenomes. http://biorxiv.org/lookup/doi/10.1101/2020.01.12.903252 (2020) doi:10.1101/2020.01.12.903252.

27. Dutilh, B. E. et al. A highly abundant bacteriophage discovered in the unknown sequences of human faecal metagenomes. Nat Commun 5, 4498 (2014).

28. Yaffe, E. & Relman, D. A. Tracking microbial evolution in the human gut using Hi-C reveals extensive horizontal gene transfer, persistence and adaptation. Nat Microbiol 5, 343–353 (2020).

29. Methé, B. A. et al. A framework for human microbiome research. Nature 486, 215–221 (2012).

30. Hendriksen, R. S. et al. Global monitoring of antimicrobial resistance based on metagenomics analyses of urban sewage. Nature Communications 10, 1124 (2019).

31. Biller, S. J. et al. Marine microbial metagenomes sampled across space and time. Scientific Data 5, 180176 (2018).

32. Galata, V., Fehlmann, T., Backes, C. & Keller, A. PLSDB: a resource of complete bacterial plasmids. Nucleic Acids Research 47, D195–D202 (2019).

33. Wallace, B. L., Bradley, J. E. & Rogolsky, M. Plasmid analyses in clinical isolates of Bacteroides fragilis and other Bacteroides species. J Clin Microbiol 14, 383–388 (1981).

34. Sóki, J. et al. Prevalence, nucleotide sequence and expression studies of two proteins of a 5.6kb, Class III, Bacteroides plasmid frequently found in clinical isolates from European countries. Plasmid 63, 86–97 (2010).

35. Moss, E. L., Maghini, D. G. & Bhatt, A. S. Complete, closed bacterial genomes from microbiomes using nanopore sequencing. Nat Biotechnol 38, 701–707 (2020).

36. Beitel, C. W. et al. Strain- and plasmid-level deconvolution of a synthetic metagenome by sequencing proximity ligation products. PeerJ 2, e415 (2014).

37. Stalder, T., Press, M. O., Sullivan, S., Liachko, I. & Top, E. M. Linking the resistome and plasmidome to the microbiome. ISME J 13, 2437–2446 (2019).

38. Cockram, C., Thierry, A., Gorlas, A., Lestini, R. & Koszul, R. Euryarchaeal genomes are folded into SMC-dependent loops and domains, but lack transcription-mediated compartmentalization. Molecular Cell 81, 459–472.e10 (2021).

39. Nissen, J. N. et al. Improved metagenome binning and assembly using deep variational autoencoders. Nat Biotechnol (2021) doi:10.1038/s41587-020-00777-4.

40. MetaHIT Consortium et al. Identification and assembly of genomes and genetic elements in complex metagenomic samples without using reference genomes. Nat Biotechnol 32, 822–828 (2014).

41. Weisberg, A. J. et al. Unexpected conservation and global transmission of agrobacterial virulence plasmids. Science 368, eaba5256 (2020).

42. Garud, N. R., Good, B. H., Hallatschek, O. & Pollard, K. S. Evolutionary dynamics of bacteria in the gut microbiome within and across hosts. PLoS Biol 17, e3000102 (2019).

43. Shi, Z. J., Dimitrov, B., Zhao, C., Nayfach, S. & Pollard, K. S. Ultra-rapid metagenotyping of the human gut microbiome. http://biorxiv.org/lookup/doi/10.1101/2020.06.12.149336 (2020) doi:10.1101/2020.06.12.149336.

44. Sakoparnig, T., Field, C. & van Nimwegen, E. Whole genome phylogenies reflect the distributions of recombination rates for many bacterial species. eLife 10, e65366 (2021).

