## Supplementary material for "Precise genotyping of circular mobile elements uncovers human associated plasmids with surprisingly recent common ancestors": Methods

### Graph theory

**Assembly graph.** A *contig* $x$ is a sequence of nucleotides from the set $\{A, G, C, T\}$, and is associated with a *head vertex* $v_x^{head}$ (5' contig end), a *tail vertex* $v_x^{tail}$ (3' contig end), and an *internal directed edge* $e(x) = (v_x^{head}, v_x^{tail})$. A pair of contigs $x, y$ is associated with an *external directed edge* $e(x, y) = (v_x^{tail}, v_y^{head})$. A contig set is called *non-redundant* if for every contig $x$ in the set the reverse complement of $x$, denoted by $x'$ is not in the set. Let $X$ be a non-redundant contig set. We denote all vertices of $X$ by $V = \bigcup_{x \in X}\{v_x^{head}, v_{x'}^{head}, v_x^{tail}, v_{x'}^{tail}\}$ and all edges of $X$ by $E = \bigcup_{x \in X}\{e(x), e(x')\} \cup \bigcup_{x,y \in X}\{e(x, y), e(x', y), e(x, y'), e(x', y')\}$. The *directed assembly graph* of the contig set $X$ is then defined to be $G = (V, E)$.

**Circuits and cycles.** Let $\bar{x} = (x_0, x_1, \dots, x_{n-1})$ be a sequence of directed contigs in $X$, or formally it holds that $(x_i \in X) \vee (x_i' \in X)$ for $i = 0, \dots, n-1$. The *circuit* $p(\bar{x})$ is the path in $G$ that traverses the graph through the vertices $V(p(\bar{x})) = (v_{x_0}^{head}, v_{x_0}^{tail}, v_{x_1}^{head}, v_{x_1}^{tail}, \dots, v_{x_{n-1}}^{head}, v_{x_{n-1}}^{tail}, v_{x_0}^{head})$, and with matching edges $E(p(\bar{x})) = (e(x_0), e(x_0, x_1), e(x_1), \dots, e(x_{n-1}, x_0))$. We note that the same circuit $p(\bar{x})$ can be represented in $n$ alternative ways by selecting a different starting contig. The circuit $p(\bar{x})$ is called a *cycle* if it does not traverse a contig or its reverse complement more than once, or formally $max_{x \in X}|\{0 \le i < n: (x_i = x) \vee (x_i = x')\}| \le 1$. The reverse complement of $p(\bar{x})$, denoted by $p(\bar{x})'$, is the circuit associated with the contig sequence $(x_{n-1}', \dots, x_0')$.

**Cycle and edge coverage.** We denote the collection of all circuits in the graph $G$ by $C$. Note that by definition there is a one-to-one relationship between $C$ and all possible circular genomes composed of oriented contigs selected from $X$. The *genome coverage* $H: C \to \mathbb{R}_{\ge 0}$ is a latent function that satisfies $H(c) = H(c')$. This condition is required since the two strands of a molecule have the same coverage by definition. $H(c)$ represents the coverages of the circular genome associated with the circuit $c$. The *edge multiplicity* $m_c(e) \ge 0$ of an edge $e \in E$ is the number of times the edge is traversed in the circuit $c$, or formally $m_c(e) = |\{x \in E(c): x = e\}|$. The *edge coverage* $W: E \to \mathbb{R}_{\ge 0}$ is the weighted sum of the circuit coverages as they traverse edges, or formally, $W(e) = \sum_{c \in C} m_c(e) \cdot H(c)$. The *observed assembly graph* over $H$, denoted by $G_{>0} = (V, E_{>0})$ and with $E_{>0} = \{e \in E: W(e) > 0\}$, is $G$ restricted to edges with positive coverages. We define $R = \{c \in C: H(c) > 0\}$. We refer to the circular genomes associated with circuits for which $H$ is positive as the underlying *genome configuration* of the community.

**Assessing genome coverages using edge coverages.** While $H$ and its induced genome configuration are latent, $W$ is an observed variable. We first assume that $W$ is known and use it to define dominant cycles. In the implementation section below, we describe how we estimate $W$ by mapping paired reads back onto the assembly. The underlying genome configuration of an evolving and complex microbial population may include the repetition of genetic sequences within the same genome or between different genomes. Accordingly, $H$ can have a positive value for partially overlapping circuits. While we allow $H$ to have a positive value on any circuit, we focus on characterizing $H$ solely over graph cycles. We do this by defining two latent and two observable variables for each cycle as follows.

**Cycle multiplicity.** Let $\bar{x} = (x_i)_{i=0}^{n_x-1}$ be the contigs of a cycle $c = p(\bar{x})$, let $\bar{y} = (y_i)_{i=0}^{n_y-1}$ be the contigs of a circuit $t = p(\bar{y})$, and let $m \geq 0$ be a non-negative natural number. We say the $c$ has a multiplicity of $m$ in $t$ if there exists $0 \leq s_x < n_x$ and $0 \leq s_y < n_y$ such that $x_{(s_x+i)(mod\ n_x)} = y_{s_y+i}$ for $i = 0, \dots, (m-1) \cdot n_x$. In words, $c$ has a multiplicity of $m$ in $t$ if there exists a subset of $t$ that traverses along the edges of $c$ while visiting one of the contigs of $c$ at least $m$ times before leaving the cycle. Note that by definition every cycle $c$ has a multiplicity of $0$ in any circuit $t$. We define the *cycle-circuit multiplicity* $\mu_t(c)$ to be the maximal $m$ for which $c$ has a multiplicity of $m$ in $t$, and define the *cycle multiplicity* $\mu(c) = \max_{t \in R} \mu_t(c)$. In words, $\mu(c)$ is equal to the maximal *cycle-circuit* multiplicity involving $c$ for which the circuit $t$ satisfies $H(t) > 0$.

**Real and phantom cycles.** Cycles are ambiguous in the sense that the precise number of cycle turns involved in producing an associated circular genome (i.e., tandem repeats) cannot be inferred from the graph. To address this inherent ambiguity, we create a proxy for the genome coverage $H$ using a generalized genome coverage $\psi$ defined as follows: For a cycle $c = p((x_0, x_1, \dots, x_{n-1}))$, we use $c^m$ to denote the circuit produced by $m \geq 1$ loops along the contigs that define the cycle, or formally $c^m = p((y_0, y_1, \dots, y_{m \cdot n-1}))$ with $y_i = x_{i\ mod\ n}$ for $i = 0, \dots, m \cdot n - 1$. We define the *generalized genome coverage* of a cycle $c$ to be $\psi(c) = \sum_{m \in \mathbb{N}} H(c^m)$, and call a cycle $c$ for which $\psi(c) > 0$ a *real cycle* and otherwise we call it a *phantom cycle*.

**Dominant cycles.** Our goal is to distinguish between real and phantom cycles in the face of an unknown and potentially complex background of genomes. This is a non-trivial task, as the assembly graphs of community-based genome configurations may be riddled with phantom cycles. For example, the mere presence of $n$ instances of an integrated repeat element result in

$n$ phantom cycles. We address this by defining dominant graph cycles as follows. Let $c$ be a cycle in the graph. Given an edge $e = (v, u)$ we say $e$ is an *outgoing edge* of $c$ if $v \in V(c)$ and $e \notin E(c)$, and denote by $E_{\text{out}}(c)$ the set of outgoing edges. Similarly, we say $e$ is an *ingoing edge* of $c$ if $u \in V(c)$ and $e \notin E(c)$, and denote by $E_{\text{in}}(c)$ the set of ingoing edges. We define the *bottleneck coverage* $\sigma(c) = \min_{e \in E(c)} W(e)$, the *outgoing coverage* $\tau_{out}(c) = \sum_{e \in E_{\text{out}}(c)} W(e)$ and the *ingoing coverage* $\tau_{in}(c) = \sum_{e \in E_{\text{in}}(c)} W(e)$. We define the *external coverage* $\tau(c) = (\tau_{out}(c) + \tau_{in}(c))/2$ and call the cycle $c$ a *dominant cycle* if $\sigma(c) > \tau(c)$. We also define the *cycle score* $s(c) = \frac{\sigma(c)}{\tau(c)}$.

**Lower bound on generalized genome coverage.** The cycle multiplicity $\mu$ and generalized genome coverage $\psi$ are two latent variables of graph cycles. The following theorem establishes an association between them as a function of the two observable cycle variables $\sigma$ and $\tau$.

**Theorem 1**. Let $X$ be a non-redundant contig set, let $G$ be the assembly graph of $X$, let $H$ be a latent genome coverage. For any cycle $c$ in $G$ it holds that $\sigma(c) - \mu(c) \cdot \tau(c) \leq \psi(c)$.

See formal proof in **Supp. Note 1**. In a nutshell, the proof is based on the fact that any circuit other than the cycle itself that contributes to the bottleneck coverage must also contribute to the external coverage on its way into and out of the cycle. The cycle multiplicity accounts for the increased contribution to the bottleneck of circuits that make multiple turns within the cycle before exiting. A simple corollary of this inequality is that if a cycle is a phantom cycle (i.e., $\psi(c) = 0$), then $\mu(c) \geq s(c)$. In other words, the multiplicity of a phantom cycle must be greater than or equal to its score.

### Datasets

**Metagenomic and reference datasets.** Reference genome sequences for 56 plasmids, 50 phage and 100 microbial host genomes were downloaded from NCBI (**Supplementary Table 1**). Shotgun data for the two focal subjects were downloaded from the SRA database, accession numbers SRR8187104 (gut sample #1) and SRR8186424 (gut sample #2). The 30 metagenomic datasets used in this work were published in studies involving human stool samples from healthy subjects[1], a marine environment[2], and wastewater[3]. For each of the 3 environments, we chose the top ten samples with the highest read count from unique geographic locations or subjects (**Supplementary Table 2**).

**Simulated plasmid and phage configurations.** A *genome configuration* is composed of a set of circular genomes with associated x-coverage values. To generate a plasmid configuration, a

50kb circular random sequence ('central allele') was generated, and n=8 circular variants were subsequently generated by introducing for each variant a single genome rearrangement event on the sequence of the central allele. The rearrangement events were selected in equal probability among insertions, inversions, and deletions. An insertion involved introducing a sequence of a length chosen uniformly from the range 100bp-10kb and inserted at a random genome coordinate of the central allele. An inversion or deletion involved a random genome segment with a length that followed a uniform distribution. For a prophage configuration, a 10kb phage circular genome sequence ('central allele') and a 1mb host chromosome circular genome sequence were generated. The sequence of the phage genome was integrated at n=8 random locations along the chromosome genome. For both plasmid and phage, the central allele was assigned an x-coverage F between 5 and 100, increasing in steps of 5. For a plasmid configuration, the x-coverage of each variant was assigned a fraction of 100-F weighted by the beta distribution $Beta(\alpha = 2, \beta = 2)$. For a phage configuration the x-coverage of the host chromosome was set to $\frac{100-F}{n}$. For each central allele frequency 30 configuration replicates were used, producing a total of 1200 configurations for the plasmid and the phage scenarios.

**Chromosomal configuration.** The simulated chromosomal metagenome configuration was created using 100 reference bacterial chromosome sequences selected from diverse taxa as specified in **Supplementary Table 1**. Each reference sequence was assigned an x-coverage sampled uniformly in the range of 10x to 75x.

**Simulating sequence data.** For each specific genome configuration (either phage, plasmid or chromosomal) a *read dataset* was generated as follows. The total number of reads for a genome was computed such that the mean x-coverage matched the x-coverage specified in the configuration, taking into account the length of each read side, which was set to 150bp. A single paired read was generated by selecting a random strand and coordinate on a genome, and with a read insertion size (i.e., distance between read sides) that followed a normal distribution with a mean of 200bp and a standard deviation of 15bp.

### Implementation overview

We first give a short overview of the implementation before diving into the nitty-gritty details. We designed an algorithm that recovers all dominant cycles in an assembly graph (**Algorithm 1**). The algorithm reports only dominant cycles (Proof in **Supp. Note 2**) and does not miss dominant cycles (Proof in **Supp. Note 3**). We implemented a tool (DomCycle) that receives a metagenomic assembly and a set of paired reads mapped to the assembly as input, then outputs all dominant cycles in the assembly graph. DomCycle works by applying first a contig-level step that identifies candidate cycles from the assembly graph. In this step, the edge coverage $W$ is estimated using mapped read densities within contigs and between contigs. Then, Algorithm 1 is applied to find a set of candidate cycles. In a second nucleotide-level step, candidate cycles are scrutinized at single nucleotide resolution. The nucleotide-level step is designed to address issues that arise from assembly limitations outside the scope of the theoretical framework, such as over-assembly (concatenated contigs that are non-adjacent in some underlying genomes) and fragmentation (i.e., missing or very short contigs). In this step, candidate cycles are tested using two nucleotide-level criteria that incorporate mapped read densities along the cycles. Broadly speaking, the nucleotide-level definitions of dominant cycles adapt the theoretical definitions to the peculiarities of metagenome assemblies. Based on the previously defined cycle score $s(c)$, we define a nucleotide-level version called the *global nucleotide-level score*. It is computed by considering all reads that map on both sides to the assembly and on at least one side anywhere along the cycle. A second score, called the *local nucleotide-level score,* was designed to directly address the distribution of singleton reads for which one side did not map to the assembly due to assembly

**Algorithm 1:** Find dominant cycles in graph.

Input: Graph $G_{>0} = G(E, V, W)$

Output: Set of dominant cycles R

$R = \emptyset$;

For each edge $e_0 = (v_0, u_0)$ in $E$ do:

  $c = (e_0)$, $\mathrm{v} = v_0$, $\mathrm{u} = u_0$, $b = W(e_0)$, $\mathrm{a} = 0$, $\mathrm{is\_open} = \mathrm{true}$;

  while $\boldsymbol{a} < \boldsymbol{b}$ and $\textbf{is\_open}$ do:

    $N = \{e = (x, y) \in E : x = \mathrm{u}\}$;

    $M = \{e \in N : W(e) \geq b\}$;

    if $|\boldsymbol{M}| \neq \mathbf{1}$ then break; $\{m\} = M$;

    $\mathrm{u} = tail\_vertex(m)$;

    $\mathrm{is\_open} = \mathrm{u} \notin cycle\_vertices(c)$;

    push_back$(c, m)$;

    $\mathrm{a} = \mathrm{a} + \sum_{e \in N \setminus m} W(e)$;

  if $\boldsymbol{v} = \boldsymbol{u}$ and $\boldsymbol{a} < \boldsymbol{b}$ then insert$(R, c)$;

return $R$;

fragmentation. The local score is designed to filter out cases in which a cycle is not real but was integrated into a larger genomic context through sequences that failed to assemble. DomCycle reports dominant cycles corresponding to candidate cycles for which both the global and the local nucleotide-level scores are significantly greater than 1. Finally, to account for stochasticity in read distribution, p-values are assigned to the two nucleotide-level scores by modeling the number of reads using a binomial distribution. DomCycle reports all dominant cycles and their corresponding genomes.

### Implementation details

**Basic read processing.** All raw reads of a given dataset were processed as follows. Exact duplicate reads were removed using a custom in-house c++ program. Sequencing adapter removal and sequencing quality filtering was performed using Trimmomatic[4] (v0.38) with command parameters "LEADING:20 TRAILING:3 MAXINFO:60:0.1 -phred33". Reads mapping to the human genome were discarded from downstream analysis using DeconSeq[5] (v0.4.3, hg38 as reference).

**Genome assembly.** Paired-end reads were assembled using MEGAHIT[6] (v1.1.3) with the parameters "--merge-level 1000,0.95 --k-min 27 --k-max 77". Contigs smaller than 231bp were discarded. To map reads back onto the assembly, read sides were trimmed to include only the first 50bp of each read side, and sides were mapped separately to the assembly using BWA[7] (v0.7.12). Downstream analysis was limited to *mapped read sides* by filtering out sides for which (i) the match length did not span the entire 50bp or (ii) the mapping edit distance was greater than 1. A FASTG was created using the contig2fastg program from the megahit toolkit (v1.1.3).

**Internal edge weight.** Intra-contig reads were classified as *forward-facing, back-facing, left-facing, and right-facing reads,* based on the relationship between strands and coordinate order. For example, a read was classified as a forward-facing read if the read side mapping to the positive strand mapped to the left (i.e., closer to the 5'-end) of the read side mapping to the negative strand. The *internal inferred molecule length* (IML) of forward-facing reads was defined as the distance between the two start coordinates of the read sides. M was defined to be the median of the IML distribution. The *max read distance* (MD) was calculated by taking the 99.5$^{th}$ percentile of the distribution of IMLs of 100,000 randomly selected forward-facing reads and adding a safety margin of 200bp. The x-coverage of a contig was estimated by $\frac{RS * RL}{CL}$, where RS is the number of mapped read sides that mapped to the contig, RL is the pre-trimmed read side length, and CL is the contig length. The two internal edges associated with a contig and its reverse complement were assigned a weight equal to the estimated x-coverage of the contig.

**External edge weight.** External edge weights were calculated based on *external reads*, defined as the union of inter-contig reads and back-facing intra-contig reads. We defined for each external read the *external inferred molecule length* $\mathrm{EML} = d_1 + d_2 - k$, where $d_i$ is the distance between the start coordinate of a read side and the coordinate of the contig end that is reached if moving in the strand direction, and k is the k-mer length used for assembly (here k=77). Each external read was associated with two corresponding external edges, i.e., $e(x, y)$ and $e(y', x')$. An external read was classified as supporting an external edge (for both associated edges) if (i) $\mathrm{EML}<\mathrm{MD}$ for the read, and (ii) the mapped contig segments were not contained in the segment of (k+3) bases at the beginning or the end of the associated contig. The weight of an external edge was defined as $\frac{2 * UPR * RL}{M} * \frac{M+k}{M}$, where UPR is the number of reads supporting the edge, and RL is the pre-trimmed read side length. The last term in the formula is included to account for discarded reads that mapped to the first or last (k+3) bases of a contig. The weighted assembly graph was composed of vertices and internal edges for all contigs and all external edges that had a positive weight.

**Classifying reads that map to a candidate dominant cycle.** Candidate dominant cycles (*candidates*) were generated from the weighted assembly graph using an implementation of Algorithm 1. In the implementation, the testing of the condition $a < b$ specified on lines 4 and 12 was omitted, to allow for a nucleotide-level cycle classification as described below. For a given candidate, all reads for which at least one side mapped to the cycle were classified into one of the following groups. A read was classified as a *cycle-support read* if (i) the two read sides mapped to opposite cycle strands after the relative orientations of the cycle contigs were accounted for and (ii) either the read was an intra-contig read with IML ≤ MD or the read bridged a pair of consecutive contig ends along the cycle and EML ≤ MD. If both read sides mapped to contigs in the cycle but conditions (i) or (ii) were not satisfied, the read was classified as an *intra-nonsupport read*. If one read side mapped to a contig in the cycle and the other mapped to a contig not in the cycle, then the read was classified as an *inter read*. The remaining case, in which one read side did not pass our mapping criteria and the other side mapped to the cycle, was classified as a *singleton read.* Read sides classified as intra-nonsupport, inter, or singleton were sub-classified into intra-nonsupport-in, intra-nonsupport-out, inter-in, inter-out, singleton-in and singleton-out, based on the relative orientation of the read side which mapped to the cycle and the cycle itself. Together, for a given cycle, there was one class that supported the cycle ('cycle-support') with a matching read count of $N_{\text{support}}$ and six non-supporting read classes with matching read counts: $N_{intra}^{in}, N_{intra}^{out}, N_{inter}^{in}, N_{inter}^{out}, N_{singleton}^{in}, N_{singleton}^{out}$. The *total non-support coverage* ($T_c$) of a candidate $c$ was set to $(N_{intra}^{in} + N_{intra}^{out} + N_{inter}^{in} + N_{inter}^{out})/2$.

**Base pair level coverages.** For a cycle $p((x_0, \ldots, x_{n-1}))$, any coordinate $t$ within a contig $x_i$ was converted to the cycle coordinate $y(t, i) = t + \sum_{j=0}^{i-1}(L_j - k)$, where $L_i$ is the length of $x_i$ and k is the k-mer size used for assembly. For every read side that mapped to a contig within a cycle, a cycle strand was determined based on the mapped contig strand and the orientation of the contig within the cycle. We define the cycle nucleotides that are *covered* by a read as follows. If the read was classified as a cycle-supporting read it covered all bases between the two cycle coordinates defined by the start of each read side, while taking into account the two facing cycle strands. In the remaining cases, in which a read was classified as an intra-nonsupport, inter, or singleton, read sides contributed separately to their respective coverage profiles. In those cases, a read side covered all cycle coordinates within the segment that started on the cycle coordinate at the start of the read side and ended $M$ base-pairs away while moving along the appropriate cycle strand. All reads were traversed and the nucleotide-level coverage profile was computed for the 7 types: supporting, in-intra, out-intra, in-inter, out-inter, in-singleton, out-singleton. Each profile was a vector of the form $n_c^P[i]$ where $c$ is a cycle, P is the profile type and $i$ is the cycle coordinate.

**Cycle scores and P-values.** For each candidate $c$, we computed the base bottleneck coverage $B_c = min_{0 \le i < L(c)}(n_c^{support}[i])$, where $L(c)$ is the length of the cycle $c$. The *global nucleotide-level score* was set to $\frac{B_c}{T_c}$. To assign a p-value for the score, we use a null hypothesis that assumes $T_c$ is distributed according to the binomial distribution $B\left(n = N,\ p = \frac{B_c}{N}\right)$, where N is the total number of reads that mapped to the assembly and compute the probability of sampling a value $x \le T_c$. The local nucleotide-level score was computed as follows: For each base position $i$ in a candidate $c$, we calculated $T_c^{local}[i] = \max(n_c^{in}[i], n_c^{out}[i])$, where $n_c^{in}[i] = n_c^{in-intra}[i] + n_c^{in-inter}[i] + n_c^{in-singleton}[i]$ and $n_c^{out}[i] = n_c^{out-intra}[i] + n_c^{out-inter}[i] + n_c^{out-singleton}[i]$. The nucleotide-level *local score* was defined to be $min_{0 \le i < L(c)}(\frac{B_c^{local}[i]}{T_c^{local}[i]})$, where $B_c^{local}[i] = n_c^{support}[i]$. We computed the p-value of the null hypothesis $B_c^{local}[i_{min}] \le T_c^{local}[i_{min}]$ for the coordinate $i_{min}$ where the minimal score was achieved using a binomial distribution as for the global score. A candidate was reported as a vetted dominant cycle if the p-value was under 0.01 for both the global and local nucleotide-level scores.

### Performance evaluation

**Tool comparison.** DomCycle was compared to metaplasmidSPAdes[8] (part of SPAdes v3.14.1), Recycler[9] (v0.62), and SCAPP[10] (downloaded June 2020) on reference-based plasmids, reference-based phage, and the simulated chromosomal metagenome. Tools were run with default parameters. Recycler and SCAPP were supplied FASTGs generated from the same assemblies used as input for DomCycle, while including contigs shorter than 231bp that are discarded by DomCycle. MetaplasmidSPAdes was run with a maximum k-mer size of 77. For Recycler and SCAPP, BAM files were generated with BWA and filtered using the view command in SAMtools[11] (v1.9) with parameters "-bF 0x0800" as recommended in https://github.com/Shamir-Lab/Recycler (Nov. 2020). For each tool, we considered output fasta files for element reporting. For SCAPP, we only considered output sequences classified as 'confident' predictions. For metaplasmidSPAdes, we considered the output file "contigs.fasta" for element reporting.

**Performance of runs.** Configurations were evaluated in the context of runs, defined as the output of running a DNA dataset associated with a configuration using a specific tool. For a given genome configuration, the sequences (fasta format) of output cycles were aligned to the original genome configuration using nucmer (run with –maxmatch) and nucmer results were parsed using show-coords (run with "-L 200 -I 99.5"). For each genome in a configuration, we calculated the percent of the genome covered by each reported cycle in a run. For each reported cycle in a run, we identified the *nearest genome* defined as the genome with the maximum number of bases covered by the reported cycle. We determined that the genome sequence order of the nearest genome was preserved by a reported cycle if the following two conditions were satisfied: (i) the alignments between the reported cycle and genome occur only on one reported cycle strand and one genome strand and (ii) when traversing from the beginning of the genome to the end, every alignment to the reported cycle strictly occurs in the order of the reported cycle sequence.

**Recall and precision for reference datasets.** For reference-based runs shown in Figure 2, a run was classified as successful if one of the reported cycles had an alignment coverage greater than 90% of the reference genome, with preserved genome sequence order. Reference genome sequences of plasmids and phage were grouped into a plasmid and a phage dataset collection.

The recall of a collection was defined as the number of successful runs divided by the number of genomes in the collection. The precision was defined as the number of successful runs in the collection divided by the total number of reported cycles in the collection.

**Recall and precision for simulated datasets.** For the simulated plasmid and phage configurations shown in Figure 3, a run was classified as successful if one of the reported cycles had an alignment coverage on the central allele greater than 98%, with preserved sequence order. The recall value associated with a configuration associated with a specific central allele frequency was set to the number of successful runs divided by the number of replicates (n=30 replicates were used). Precision was defined as the number of runs aligning to one of the genomes in the configuration with coverage greater than 98% and sequence order preserved divided by the number of reported cycles. If no cycles were reported, precision was defaulted to 100%.

### Cycle characterization

**Gene annotation.** Genes were predicted on dominant cycles using prodigal[12] (v2.6.3). Translated gene predictions were aligned to UNIREF100[13] (downloaded July 2020) using diamond[14] blastp with "–sensitive" and a maximum e-value of 0.001. For a gene with multiple alignments to UNIREF100, the alignment with the greatest sequence identity was kept, where identity is defined as the alignment similarity multiplied by the fraction of the target gene that was covered by the alignment. HMMER hmmscan[15] (v3.1.b2) was used to report cycle gene alignments to the Pfam database[16] (downloaded August 2019) with a maximum e-value of 0.001.

**Cycle classification.** The names of gene hits in the UNIREF100 and Pfam databases were used to classify dominant cycle into one of the following functional categories: plasmid, phage, mobile, or undefined. A cycle was assigned to a functional category if one or more of its genes matched one of the following regular expressions, tested while accepting both lower and upper case. The *plasmid* category expressions used were: "plasmid", "conjug.*", "trb$", "Mob[A-E]$*", "Par[A-B]". The *phage* category expressions were: "capsid", "phage.*", "tail", "head", "tape", "antitermination", "virus.*", "Bacteriophage", "sipho*", "Baseplate", "T4-like.*", "myovir.*". The *mobile* category expressions were: "transpos.*","resolvase", "toxin", "antitoxin", "excision*", "integrase", "relaxase", "recombination", "segregation", "extrachromosomal", "mobilization", "partitioning". If a cycle met classification for more than one category, then the cycle was assigned to the first matched category in the list: *phage*, *plasmid*, *mobile.* If a cycle matched none of the three categories it was assigned to the *undefined* category. Circulating cycles (i.e., associated with one of the 20 clusters) were further annotated as described below.

**Cycle coverages and abundance percentiles.** We define the *adjusted median coverage* (AMC) of a dominant cycle $c$ to be $M_c - T_c$ where $M_c$ denotes the median base pair support. For each sample separately, we generated *pseudo-dominant genomes* (PDGs) as follows. We considered *candidate PDGs* as contigs that do not contribute to a dominant cycle and were larger than $2 * MD$. Profiles were computed for candidate PDGs using the same read classification procedure used for candidates. The PDG base bottleneck coverage was computed while avoiding contig sides, or formally $B'_p = min_{MD \le i < (L(c) - MD)}(n_p^{support}[i])$. The *total non-support coverage* of PDGs ($T'_p$) was also computed while avoiding the first and last MD bases on the contig. The *global nucleotide-level score* of PDGs was set to $\frac{B'_p}{T'_p}$. Similarly, the local score for a candidate PDG was computed while omitting positions that were up to MD bases from contig ends. A candidate PDG was classified as a vetted PDG if it passed the global nucleotide score test and the local score test. The AMC of a vetted PDG $p$ was set to $M'_p - T'_p$, where $M'_p$ is the median support coverage computed over contig bases between the first and last MD bases. The *abundance percentile* of a dominant cycle was set to the percentile of the AMC of the cycle within the distribution of AMC values across all PDGs in the sample. One-sided Kolmogorov-Smirnov tests were used to test whether a distribution of dominant cycle AMCs were enriched over a background of AMCs. To calculate the coverage distribution of contigs in the assembly, we calculated the median coverage for each PDG along the bases between the first and last MD bases in the contig.

### Cycle clustering

**MGE clusters.** The corresponding genomes of all dominant cycles recovered from the 32 samples described in this study were aligned in pairs using nucmer (run with "--maxmatch", part of the MUMmer v3.1 package[17]). Single nucleotide polymorphisms (SNPs) between cycle pairs were identified using show-snps (part of MUMmer). The alignment metrics for two cycles $c_i$ and $c_j$ were defined as follows. The *alignment fraction* was $F(c_i, c_j) = \left(\frac{O(c_i)}{L(c_i)} + \frac{O(c_j)}{L(c_j)}\right) / 2$, where $O(c)$ denotes the number of bases in cycle $c$ covered by the alignment and $L(c)$ denotes the length of cycle $c$. The *alignment identity* $I(c_i, c_j)$ was the average nucleotide identity (ANI) within the aligned fraction. The *weighted alignment identity* (referred to as the ANI in the main text) was set to $A(c_i, c_j) = F(c_i, c_j) * I(c_i, c_j)$, effectively counting non-aligned regions as having zero identity. The *genome distance* used to cluster cycles was $D(c_i, c_j) = 1 - A(c_i, c_j)$. Cycles were clustered based on their genome distances using a threshold of 0.05 (equal to 95% ANI) and using single linkage. A single cluster associated with the PhiX genome and all clusters with a single member were discarded from the analysis. All cycles associated with multi-member clusters were termed *circulating cycles*.

**PLSDB references**. A representative cycle for each cluster in a circulating species was aligned to PLSDB[18] using 'mash dist' (distance cutoff of 0.25) through the PLSDB webserver. Each PLSDB reference was downloaded and aligned using nucmer and genome distances were computed in pairs between all reference sequences and the genomes of circulating cycles, as described above.

**Annotation of circulating cycles.** For each cycle $c$ belonging to a multi-member cluster, we predicted genes by triplicating the cycle sequence and concatenating each copy together sequentially. We only considered genes on cycle $c$ where the gene start coordinate $x$ (on the triplicated cycle sequence) satisfied $L(c) < x \leq 2 * L(c)$. In addition to the annotation procedure described in the section "Gene annotation" above, HMMER hmmscan was used to align (with a max e-value of 0.001) genes to families of mobilization genes, secretion systems, and conjugation genes downloaded from COPLA[19]. Separately, Pfam and UNIREF gene classifications were inspected for toxin-antitoxin genes and replication genes. The complete annotations of all genes on circulating cycles are found in **Supp. table 3**. The MOB column in Table 1 was determined based on Relaxase HMM hits to the mobilization, secretion or conjugation gene families obtained from COPLA. The 'uncharacterized' column in Table 1 was determined based on the number of genes that had no HMM/Pfam hits and either had no hits in UNIREF or matched an uncharacterized or hypothetical protein. The addiction column in Table 1 was determined based on Toxin-antitoxin Pfam and UNIREF annotations and included descriptions such as "Antitoxin Phd_YefM", "YoeB-like toxin", "ParE toxin", "Antitoxin VbhA", "Antidote-toxin recognition MazE", "PemK-like, MazF-like toxin", and "RelE toxin". The replication column in Table 1 was determined based on the Pfam and UNIREF descriptions "Replication protein", "RepB family", "Initiator replication protein", "Replication factor-A C terminal domain", and "Firmicute plasmid replication protein (RepL)". The 16 clusters that had a MOB or replication gene were classified as plasmids. Clusters M6, M12, M13 and M15 were left undetermined.

**Phylogenetic trees.** The phylogeny trees in Figure 6 were generated as follows. Given a reference 'pivot' cluster member, all SNPs that separated the pivot member and other cluster members (identified by nucmer) were used to generate a multi-sequence alignment based on the reference genome of the pivot member. In case of an indel (denoted by a '.' by show-coords of nucmer) a gap was placed in the alignment. PhyML[20] was run on the alignment with default parameters and optionally marking an outgroup (see below). In the case of M1, the 3 members of M1b were marked as outgroup members when running PhyML for visualization purposes. Similarly, the *Enterobacter hormaechei* reference was marked as an outgroup for M18.

### Code availability

DomCycle was implemented in python and R and is available for download as an open-source tool at https://github.com/nshalon/DomCycle.

#### References


1. Methé BA, Nelson KE, Pop M, et al. A framework for human microbiome research. *Nature*. 2012;486(7402):215-221. doi:10.1038/nature11209

2. Biller SJ, Berube PM, Dooley K, et al. Marine microbial metagenomes sampled across space and time. *Scientific Data*. 2018;5(1):180176. doi:10.1038/sdata.2018.176

3. Hendriksen RS, Munk P, Njage P, et al. Global monitoring of antimicrobial resistance based on metagenomics analyses of urban sewage. *Nature Communications*. 2019;10(1):1124. doi:10.1038/s41467-019-08853-3

4. Bolger AM, Lohse M, Usadel B. Trimmomatic: a flexible trimmer for Illumina sequence data. *Bioinformatics*. 2014;30(15):2114-2120. doi:10.1093/bioinformatics/btu170

5. Schmieder R, Edwards R. Fast Identification and Removal of Sequence Contamination from Genomic and Metagenomic Datasets. *PLOS ONE*. 2011;6(3):e17288. doi:10.1371/journal.pone.0017288

6. Li D, Liu C-M, Luo R, Sadakane K, Lam T-W. MEGAHIT: an ultra-fast single-node solution for large and complex metagenomics assembly via succinct de Bruijn graph. *Bioinformatics*. 2015;31(10):1674-1676. doi:10.1093/bioinformatics/btv033

7. Li H, Durbin R. Fast and accurate short read alignment with Burrows–Wheeler transform. *Bioinformatics*. 2009;25(14):1754-1760. doi:10.1093/bioinformatics/btp324

8. Antipov D, Raiko M, Lapidus A, Pevzner PA. Plasmid detection and assembly in genomic and metagenomic data sets. *Genome Res*. 2019;29(6):961-968. doi:10.1101/gr.241299.118

9. Rozov R, Brown Kav A, Bogumil D, et al. Recycler: an algorithm for detecting plasmids from de novo assembly graphs. *Bioinformatics*. 2017;33(4):475-482. doi:10.1093/bioinformatics/btw651

10. Pellow D, Zorea A, Probst M, et al. *SCAPP: An Algorithm for Improved Plasmid Assembly in Metagenomes*. Bioinformatics; 2020. doi:10.1101/2020.01.12.903252

11. Li H, Handsaker B, Wysoker A, et al. The Sequence Alignment/Map format and SAMtools. *Bioinformatics*. 2009;25(16):2078-2079. doi:10.1093/bioinformatics/btp352

12. Hyatt D, Chen G-L, LoCascio PF, Land ML, Larimer FW, Hauser LJ. Prodigal: prokaryotic gene recognition and translation initiation site identification. *BMC Bioinformatics*. 2010;11:119. doi:10.1186/1471-2105-11-119

13. Suzek BE, Huang H, McGarvey P, Mazumder R, Wu CH. UniRef: comprehensive and non-redundant UniProt reference clusters. *Bioinformatics*. 2007;23(10):1282-1288. doi:10.1093/bioinformatics/btm098

14. Buchfink B, Xie C, Huson DH. Fast and sensitive protein alignment using DIAMOND. *Nat Methods*. 2015;12(1):59-60. doi:10.1038/nmeth.3176

15. HMMER. Accessed November 7, 2020. http://hmmer.org/

16. Finn RD, Bateman A, Clements J, et al. Pfam: the protein families database. *Nucleic Acids Res*. 2014;42(Database issue):D222-D230. doi:10.1093/nar/gkt1223

17. Kurtz S, Phillippy A, Delcher AL, et al. Versatile and open software for comparing large genomes. *Genome Biology*. 2004;5(2):R12. doi:10.1186/gb-2004-5-2-r12

18. Galata V, Fehlmann T, Backes C, Keller A. PLSDB: a resource of complete bacterial plasmids. *Nucleic Acids Research*. 2019;47(D1):D195-D202. doi:10.1093/nar/gky1050

19. COPLA. Accessed February 2, 2021. https://castillo.dicom.unican.es/copla_about/

20. Guindon S, Dufayard J-F, Lefort V, Anisimova M, Hordijk W, Gascuel O. New Algorithms and Methods to Estimate Maximum-Likelihood Phylogenies: Assessing the Performance of PhyML 3.0. *Systematic Biology*. 2010;59(3):307-321. doi:10.1093/sysbio/syq010
