## Supplementary Notes 1-4 for "Precise genotyping of circular mobile elements uncovers human associated plasmids with surprisingly recent common ancestors"

### Supplementary Note 1

**Theorem 1**. Let $X$ be a non-redundant contig set, let $G$ be the assembly graph of $X$, let $H$ be a latent circuit *coverage* function, and let $W$ be the induced edge *coverage* function of $H$. For any cycle $c$ in $G$, it holds that $\sigma(c) - \mu(c) \cdot \tau(c) \leq \psi(c)$.

**Proof.** Let $c$ be a cycle and let $t = p(\bar{x})$ be a circuit over the contigs $\bar{x} = (x_i)_{i=0}^{n-1}$ that satisfies $H(t) > 0$. We denote by $\phi_t(c)$ the number of times the circuit $t$ crosses into the cycle $c$, or formally $\phi_t(c) = \sum_{e \in E_{\text{in}}(c)} m_t(e)$. Since every entry of the circuit into the cycle has a matching exit event it holds that $\phi_t(c) = \sum_{e \in E_{\text{out}}(c)} m_t(e)$. We first express $\tau_{in}(c)$ as a function of $\phi_t(c)$:

$$\tau_{in}(c) = \sum_{e \in E_{\text{in}}(c)} W(e) = \sum_{e \in E_{\text{in}}(c)} \sum_{t \in R} m_t(e) \cdot H(t) = \sum_{t \in R} H(t) \sum_{e \in E_{\text{in}}(c)} m_t(e) = \sum_{t \in R} H(t) \cdot \phi_t(c)$$

Similarly, $\tau_{out}(c)$ also equals $\sum_{t \in C} H(t) \cdot \phi_t(c)$ and therefore:

$$\tau(c) = \sum_{t \in R} H(t) \cdot \phi_t(c) \qquad (1)$$

The set of circuits $R$ can be expressed as a disjoint union of (1) the circuits $R_{out}$ that do not cross into the cycle $c$, (2) the circuits $R_{in}$ that do not cross out of the cycle $c$, and (3) the circuits $R_{cross}$ that cross into the cycle $c$ once or more.

Let $e_0 = \underset{e \in E(c)}{\operatorname{argmin}} W(e)$, be an edge in the cycle that gets a minimal coverage value. It holds that:

$$\sigma(c) = W(e_0) = \sum_{t \in R} m_t(e_0) \cdot H(t)$$

$$= \sum_{t \in R_{out}} m_t(e_0) \cdot H(t) + \sum_{t \in R_{cross}} m_t(e_0) \cdot H(t) + \sum_{t \in R_{in}} m_t(e_0) \cdot H(t)$$

Since $\forall t \in R_{out}: (m_t(e_0) = 0)$ and $\psi(c) = \sum_{t \in R_{in}} m_t(e_0) \cdot H(t)$ (by definition) we can simplify:

$$\sigma(c) = \sum_{t \in R_{cross}} m_t(e_0) \cdot H(t) + \psi(c)$$

By definition, any circuit $t \in R_{cross}$ enters the cycle $c$ precisely $\phi_t(c)$ times, and traverses the edge $e_0$ up to $\mu(c)$ times on each visit. Therefore, it holds that $m_t(e_0) \leq \mu(c) \cdot \phi_t(c)$. We substitute $m_t(e_0)$ to get the inequality:

$$\sigma(c) \leq \sum_{t \in R_{cross}} \mu(c) \cdot \phi_t(c) \cdot H(t) + \psi(c)$$

Reorganizing the equation, we get:

$$\sigma(c) - \sum_{t \in R_{cross}} \mu(c) \cdot \phi_t(c) \cdot H(t) \leq \psi(c)$$

By assigning equation 1 we get:

$\sigma(c) - \mu(c) \cdot \tau(c) \leq \psi(c)$ ■

#### Supplementary Note 2

**Proposition:** All cycles returned by Algorithm 1 are dominant.

**Proof.** Assume $c$ is added to the result set $R$ in line 12 of Algorithm 1. It holds that $c$ is a cycle since $v = u$, i.e., the path is closed. The variable $b$ is equal to the weight $e_0$ (line 3) and verified to be equal or smaller than the weight of all edges in $c$ (line 6), and therefore by definition $b = \sigma(c)$. The variable $a$ equals the $\tau_{out}(c)$ by construction due to line 11. Therefore, it holds that $\tau_{out}(c) = a < b = \sigma(c)$. According to Supplementary Note 1 we know that $\tau_{out}(c) = \tau_{in}(c)$ and thus $\tau(c) < \sigma(c)$ ■

#### Supplementary Note 3

**Proposition:** Algorithm 1 returns all dominant cycles.

**Proof.** Assume $c$ is a dominant cycle in the graph. Let $e \in E(c)$ satisfy $W(e) = \sigma(c)$. Let us focus on the two algorithm loop attempts (in line 2) that start from $e$ and $e'$. Suppose for the sake of contradiction that both runs fail and therefore $c$ or $c'$ are not reported by the algorithm. Therefore, the algorithm leaves the cycle prematurely for both attempts and there exists two edges $e_{in} \in E_{\text{in}}(c)$ and $e_{out} \in E_{\text{out}}(c)$ such that $W(e_{in}) \geq \sigma(c)$ and $W(e_{out}) \geq \sigma(c)$. Averaging the two inequalities we get $(W(e_{in}) + W(e_{in}))/2 \geq \sigma(c)$. By the definition of the external coverage, we have $\tau(c) = (\tau_{out}(c) + \tau_{in}(c))/2 \geq (W(e_{in}) + W(e_{in}))/2$, and therefore $\tau(c) \geq \sigma(c)$, in contradiction to the fact that the cycle is dominant. Thus $c$ or $c'$ are reported by Algorithm 1 ■

#### Supplementary Note 4

We estimate a lower bound on the expected time to coalescence of M1a as follows. We estimate the expected number of generations until coalescence $T_c$ based on the equation $\pi = 2T_c\mu$, where $\pi$ is the nucleotide diversity (average fraction of sites that differ between pairs of genomes) and $\mu$ is the mutation rate (mutations/bp/generation). We denote by $g$ the number of generations per year which allows us to express the time to coalescence in years as $Y = \frac{T_c}{g} = \frac{\pi}{2g\mu}$.

We estimate $g\mu$ as follows: According to the two references we identified for M1, the bacterial hosts of plasmid M1a are *Bacteroides xylanisolvens* (M1a) and *Bacteroides fragilis* (M1b). We use the mutation accumulation rate of *B. fragilis* in natural conditions that was estimated at 0.9 mutations/genome/year[1]. Dividing by the size of the *B. fragilis* genome (5.295Mb) we get $g\mu = 1.699717e-07$ mutations/bp/year.

We estimate $\pi$ as follows: Using the pairwise nucmer alignments (using show-coords) of the 8 M1a members, we found that pairs are separated on average by 1.75 SNPs. Dividing by the size of M1 (4148bp) we get $\pi = 0.0004218901$.

Plugging in the estimates of $\mathrm{g}\mu$ and $\pi$, we get $Y = \frac{\pi}{2\mathrm{g}\mu} = \frac{0.0004218901}{2*1.699717\mathrm{e}-07} = 1241$ years.

Noting that the plasmid in gut #2 is an outlier in terms of the number of differentiating SNPs, we computed the same estimate of years to coalescence without that plasmid. In that case we have only 0.85 SNPs between pairs on average, which results in $\pi = 0.000204918,$ and $Y = 602$ years. One caveat worth mentioning is that this analysis is typically performed using neutral mutations (such as intra-genic synonymous mutations). Due to the small number of polymorphic sites, we use here all mutations (both in the estimation of $\mu$ and of $\pi$).

#### REFERENCES

1. Zhao S, Lieberman TD, Poyet M, et al. Adaptive Evolution within Gut Microbiomes of Healthy People. *Cell Host & Microbe*. 2019;25(5):656-667.e8. doi:10.1016/j.chom.2019.03.007
